# Introgression across narrow contact zones shapes the genomic landscape of phylogenetic variation in an African bird clade

**DOI:** 10.1101/2024.08.13.607717

**Authors:** Loïs Rancilhac, Stacey G. de Souza, Sifiso M. Lukhele, Matteo Sebastianelli, Bridget O. Ogolowa, Michaella Moysi, Christos Nikiforou, Tsyon Asfaw, Colleen T. Downs, Alan Brelsford, Bridgett M. vonHoldt, Alexander N.G. Kirschel

**Affiliations:** Department of Biological Sciences, University of Cyprus, PO Box 20537 Nicosia 1678, Cyprus; Department of Wildlife and Protected Area Management, Wondo Genet College of Forestry and Natural Resources, Hawassa University, Ethiopia; Evolutionary Ecology Group, Biology, University of Antwerp, Antwerp, Belgium; Centre for Functional Biodiversity, School of Life Sciences, University of KwaZulu-Natal, Pietermaritzburg 3209, South Africa; Department of Evolution, Ecology and Organismal Biology, University of California Riverside, Riverside, CA 92521, USA; Department of Ecology & Evolutionary Biology, Princeton University, Princeton, NJ 08544, USA

**Keywords:** Phylogenomics, Gene-tree heterogeneity, Recombination, Hybrid zones, Speciation, Birds

## Abstract

Genomic analyses of hybrid zones provide excellent opportunities to investigate the consequences of introgression in nature. In combination with phylogenomics analyses, hybrid zone studies may illuminate the role of ancient and contemporary gene flow in shaping variation of phylogenetic signals across the genome, but this avenue has not been explored yet. We combined phylogenomic and geographic cline analyses in a *Pogoniulus* tinkerbird clade to determine whether contemporary introgression through hybrid zones contributes to gene-tree heterogeneity across the species ranges. We found diverse phylogenetic signals across the genome with the most common topologies supporting monophyly among taxa connected by secondary contact zones. Remarkably, these systematic conflicts were also recovered when selecting only individuals from each taxon’s core range. Using analyses of derived allele sharing and “recombination aware” phylogenomics, we found that introgression shapes gene-tree heterogeneity, and the species tree most likely supports monophyletic red-fronted tinkerbirds, as recovered in previous reconstructions based on mitochondrial DNA. Furthermore, by fitting geographic clines across two secondary contact zones, we found that introgression rates were lower in genomic regions supporting the putative species tree compared to those supporting the two taxa in contact as monophyletic. This demonstrates that introgression through narrow contact zones shapes gene-tree heterogeneity even in allopatric populations. Finally, we did not find evidence that mitochondria-interacting nuclear genes acted as barrier loci. Our results show that species can withstand important amounts of introgression while maintaining their phenotypic integrity and ecological separation, raising questions regarding the genomic architecture of adaptation and barriers to gene flow.

## Introduction

Phylogenetic relationships vary across the genome (Degnan and Rosenberg 2009), and whole genome sequence studies have shown that more data may not provide a clearly conclusive phylogeny (Jeffroy et al. 2006; Bravo et al. 2019). Despite extensive efforts in the context of the multispecies coalescent framework, incomplete lineage sorting (ILS) and introgression may result in topological discordance across the genome, and thus remain a major challenge in the reconstruction of species trees (Scornavacca and Galtier 2017). Phylogenetic reconstructions are further complicated by genome-wide heterogeneity in recombination rate, selection, and demographic events (Pease and Hahn 2013; Schumer et al. 2018; Li et al. 2019; Musher et al. 2024; Thom et al. 2024). Although reconstructing the species tree remains a core goal of phylogenetics, studying patterns of gene-tree heterogeneity can illuminate the genetic architecture of reproductive barriers (Stankowski et al. 2024), and reveal the processes underlying repeated evolution of phenotypic traits (Alaei Kakhki et al. 2023; Rancilhac et al. 2024). Investigating gene-tree heterogeneity can thus yield insights into the factors underlying genome evolution in wild populations, and the consequences of processes such as ILS and introgression on the evolution of phenotypic traits (e.g., Feng et al., 2022).

In sexually reproducing organisms, gene tree heterogeneity chiefly results from two processes (ILS and introgression). The former is a neutral process resulting from the retention of standing variation in non-sister lineages, which depends on the effective population sizes (*Ne*) and divergence times (Degnan and Rosenberg 2009). Introgression, the transfer of genetic material across lineages through interbreeding, is more difficult to predict because its occurrence depends on the lineages’ biogeographic history to include secondary contact for gene flow to occur (Burbrink and Gehara 2018) and that admixtures are reproductively viable. The temporal persistence of introgressed variants depends on both neutral sorting processes and selection (against or favoring introgressed variants) (Edelman and Mallet 2021). Reproductive isolation increases with divergence between lineages, thus reducing gene tree heterogeneity because of introgression at deeper evolutionary timescales. Yet introgression has been documented across a range of branch divergence levels in the Tree of Life, from incipient species to distinct genera (Mallet et al. 2016). Several studies have highlighted the role of introgression in generating novel traits, especially in the context of adaptive radiations (Pease et al. 2016; Svardal et al. 2020; Meier et al. 2023). Consequently, introgression is now recognized as a fundamental and common phenomenon, but the biogeographic context in which it occurs is rarely well understood (Vanderpool et al. 2020).

Gene-tree heterogeneity can be a legacy of introgression occurring at early stages of speciation where introgressed alleles persist in present-day populations even in the absence of gene flow. This phenomenon of “ancestral introgression” is well exemplified by reported introgression between strictly allopatric species (e.g., Ferreira et al. 2021) or distantly related lineages with now-complete reproductive isolation (e.g., Rancilhac et al. 2021; Suvorov et al. 2022). By contrast, the impact of contemporary introgression on patterns of gene-tree heterogeneity in parapatric species has garnered little attention (but see Thom et al. 2018). Because gene flow is not homogeneous across space, we expect that the effect of contemporary gene flow on gene-tree genealogies would depend on the geographical sampling considered. More specifically, we hypothesize that the populations furthest away from hybrid zones are less likely to be introgressed, and their relationships should thus reflect the species tree. However, neutral genetic variants may be introgressed far from the contact zone (Payseur 2010; Cruzan et al. 2021). How neutral introgression across hybrid zones can affect the genomic landscape of genealogical variation, and the geographic scale of this phenomenon, has not been documented. Combining hybrid zone analysis with patterns of gene-tree heterogeneity at the scale of the whole genome may provide a powerful tool to address this question. Indeed, geographic clines fitted to allele frequencies at single-nucleotide polymorphisms across the genome provide estimates of local rates of introgression (Dufresnes et al. 2021), which can then be compared to local phylogenetic signals.

In this study, we investigated the contribution of ongoing gene flow to genome-wide phylogenetic conflicts in the red- and yellow-fronted tinkerbirds (*Pogoniulus pusillus-chrysoconus* species group). This group of African barbets is currently classified into two species (Kirschel et al. 2021), the red-fronted tinkerbird (*P. pusillus*, hereafter RFT) and yellow-fronted tinkerbird (*P. chrysoconus*, hereafter YFT), each of which are further divided into three subspecies (RFT: *P. p. affinis, pusillus,* and *uropygialis*; YFT: *P. c. chrysoconus, extoni* and *xanthostictus*) with limited or no range overlap between subspecies (Fig. 1). RFT has a disjunct distribution, with two subspecies in Eastern African (*P. p. affinis and P. p. uropygialis*) and the nominate form in Southern Africa (*P. p. pusillus*). YFT occupies most of central and western Africa, where one lineage found in the northern savanna region contains *P. c. chrysoconus* and *P. c. xanthostictus* with *P. c. extoni* further south (Nwankwo et al. 2019). The YFT subspecies do not form a monophyletic group in previous mtDNA-based reconstructions (Nwankwo et al. 2019; Kirschel et al. 2021). The distribution of taxa in Ethiopia is especially complicated where contact zones exist between *P. c. xanthostictus*, *P. p. uropygialis*, and *P. p. affinis* (Kirschel et al. 2021). As a result, RFT and YFT form an intricate pattern of replacement and contact zones across Africa. Assessment of the contact zone between *P. c. extoni* and *P. p. pusillus* in southern Africa based on microsatellites (Nwankwo et al. 2019) and whole genome sequences (Sebastianelli et al. 2024) reported rampant introgression between the two species. Thus, RFT and YFT provide an ideal system to investigate the contribution of contemporary introgression to gene-tree heterogeneity. We collected whole genome resequencing (WGS) data of tinkerbirds across Africa to investigate their phylogenetic relationships and patterns of gene-tree heterogeneity in the light of secondary contact zones. We further supported the WGS data with double-digest restriction-site associated DNA sequencing (ddRADseq) across two contact zones to quantify introgression and determine whether support for specific topologies is influenced by high rates of introgression. We then leveraged our understanding of introgression and gene-tree heterogeneity patterns to investigate the role of genes associated with forecrown coloration, and mitochondria interacting nuclear genes, as barriers to gene flow.

**Figure 1.**
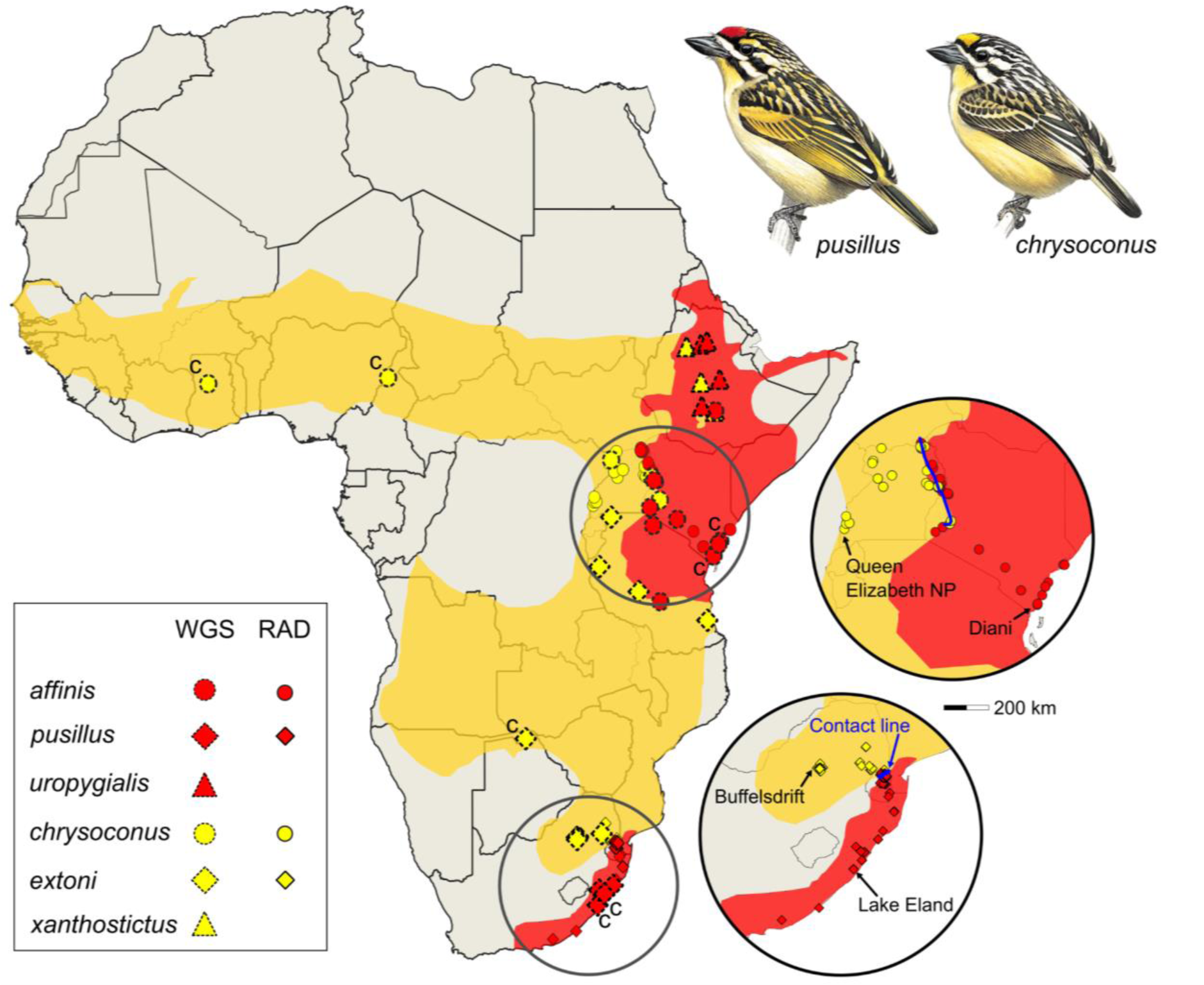
Map of the samples used in this study with the approximate ranges of P. chrysoconus (yellow) and P. pusillus (red). Markers show sampling locations for the WGS dataset (large symbols with dotted contour) and the RADseq dataset (small symbols with solid contour). Core range WGS samples are indicated with a “c”. Insets: details of the contact zones between P. c. chrysoconus and P. p. affinis in Uganda and Kenya (top) and between P. c. extoni and P. p. pusillus in Southern Africa (bottom). Markers in the insets detail the sampling used for geographic clines inference (RADseq data only), and solid blue lines show the location of the contact zones. The most distant from the contact zones, used to identify differentially fixed SNPs, are indicated with arrows (Note: for pusillus, two localities further away from the contact zone were not included in the single-SNP analysis because they were represented by single samples). Tinkerbird paintings courtesy of del Hoyo et al. (2020).

## Methods

### Study design and genomic DNA extractions

We investigated the genomic landscape of phylogenetic variation in the *Pogoniulus chrysoconus*-*pusillus* complex, and the biogeographic processes contributing to phylogenetic discordances. We generated a whole genome resequencing dataset (WGS dataset) of 43 samples covering all taxa in the species complex and two genomes from outgroup species. This dataset included representatives from both strictly allopatric localities and localities adjacent to contact zones. We additionally collected double-digest Restriction enzyme-Associated DNA sequencing data (ddRADseq datasets) across two contact zones with the expectation that collected samples represent a range of admixture along geographic transitions: the first contact zone is in Uganda and Kenya for *P. c. chrysoconus* and *P. p. affinis* (98 samples obtained during fieldwork described in Sebastianelli et al. 2022 and Kirschel & Sebastianelli 2022). The second contact zone is in Southern Africa, between *P. c. extoni* and *P. p. pusillus* (523 samples), for which samples were collected and ddRADseq performed previously (see Nwankwo et al. 2019, Kirschel et al. 2020, Sebastianelli et al. 2024). A spreadsheet with metadata for all ddRADseq samples is available as supplementary information (Appendix 1).

We extracted genomic DNA from blood samples for generating the WGS and ddRADseq datasets using a Qiagen DNeasy blood and tissue kit following the manufacturer’s protocols (Qiagen, Germany). We quantified DNA on a Qubit 2.0 fluorometer (Thermo Fisher Scientific Inc).

### Whole Genome resequencing dataset

The WGS dataset included three RFT subspecies (*P. p. affinis*: N=10, *P. p. pusillus*: N=7, *P. p. uropygialis*: N=4) and three YFT subspecies (*P. c. chrysoconus*: N=7, *P. c. extoni*: N=10, *P. c. xanthostictus*: N=5) resequenced at an expected depth of approximately 15-fold coverage (Fig. 1, Table S1). We also included a sample from each of the closely related species *P. atroflavus* and *P. bilineatus* as outgroups. Whole genome libraries were generated using the NEBNext DNA Library Prep Kit following the manufacturer’s recommendations and purified libraries using the AMPure XP system. Each library was quality checked for size distribution using an Agilent 2100 Bioanalyzer, and quantified using real-time PCR, with paired-end 2×150nt sequence data collected on an Illumina Novaseq6000 platform.

We trimmed remnant adapters and clipped low quality nucleotides (--quality-cutoff 20) using cutadapt v4.0 (Martin 2011) and discarded reads shorter than 20bp (--minimum-length 20). We next aligned the processed sequence reads to the *Pogoniulus pusillus* reference genome (GCA_015220805.1) using bwa-mem v0.7.1 (Li and Durbin 2009) and subsequently used Samtools v1.6 (Danecek et al. 2021) to exclude reads with low mapping qualities (--min-MQ 20), sort the reads by coordinate, and convert to BAM formats. We removed duplicate reads with the *MarkDuplicates* tool v2.23.1 in Picard (Broad Institute 2019). We discovered variants after base quality score recalibration in GATK v4.2.3 (McKenna et al. 2010) following best practices. We used vcftools v0.1.16 (Danecek et al. 2011) to retain only biallelic sites with a minor allele frequency (MAF) ≥0.05, a depth of coverage between 4X and 50X, variants with less than 60% missing data across samples, and quality scores above 20 (--remove-indels --maf 0.05 --minDP 4 --maxDP 50 --max-missing 0.4 --minQ 20 –minGQ 20).

We inferred the sex of the samples by calculating the ratio of average autosomal depth over average chromosome Z depth, with the expectation that females are half the coverage relative to males (Kirschel et al. 2020; Sebastianelli et al. 2024). In practice, we estimated mean individual depths for chromosome 1 and chromosome Z using vcftools v0.1.16, and calculated ratios using a custom R script. Next, we separated male and female individuals in different vcf files, and replaced all heterozygous genotypes from the female vcf with missing data using a custom bash script. Finally, we merged back the males and females using bcftools v1.9 (Danecek et al., 2021), and filtered the sites to remove positions with more than 60% missing genotypes.

### Restriction-site Associated DNA sequencing

We generated ddRAD-seq libraries using the *SbfI* and *MseI* restriction enzymes following the protocol described in Brelsford et al. (2016), which includes elements of two previously published protocols (Parchman et al. 2012; Peterson et al. 2012). We used 24 barcoded adapters and 8 indexed primers which allowed us to pool 192 samples with equimolar concentrations. We collected paired-end 2×150nt sequencing reads on an Illumina HiSeq X Ten platform by Novogene Corporation Inc.

We cleaned sequencing data with subsequent variant discovery for each of the two datasets independently (dataset 1: Uganda / Kenya; dataset 2: Southern Africa). We demultiplexed each pool using the *process_radtags* module in Stacks v2.62 (Rochette et al. 2019), discarded reads with uncalled bases (--clean). We aligned the paired-end reads to the *P. pusillus* reference genome (GCA_015220805.1) using bwa-mem v0.7.1. We filtered the resulting SAM files to remove reads with low mapping qualities (--min-MQ 20) using Samtools v1.6, and then we sorted the reads by their genomic coordinates before converting them to BAM formats. We cataloged SNP loci using the *gstacks* module and increased the alpha significance thresholds for both SNP discovery (--var-alpha 0.01) and genotype calling (--gt-alpha 0.01).

### Phylogenomic analyses

We used the WGS dataset to reconstruct the phylogenetic relationships of the red- and yellow-fronted tinkerbirds. First, we inferred Neighbor-Joining (NJ) phylogenetic trees in 500 SNP non-overlapping windows for the autosomes and the Z chromosome using the TopoWindows R script (https://github.com/rancilhac/TopoWindows/tree/main) with a Jukes-Cantor 69 model (Jukes and Cantor 1969). We then used the NJ trees to infer a phylogeny while accounting for ILS with the software ASTRAL. Because ASTRAL assumes free recombination across loci, and no within-loci recombination, we filtered the NJ trees to retain only those corresponding to windows separated by ≥10 Kb but ≤50 Kb in size, and used them as input for ASTRAL-II v5.7.8 (Mirarab et al. 2014). We ran this analysis on autosomal and Z chromosome windows separately (birds have a ZW sex chromosomes system; Irwin 2018).

Next, we investigated patterns of topological variation using a topology weighting framework, which quantifies the support of a given gene tree *t* to the possible species trees by calculating the proportion of subtrees in *t*, obtained by sub-sampling a single tip per species, that supports each species tree. Loci under linkage disequilibrium can be included in this analysis, but within-locus recombination should be avoided. Thus, we retained all NJ trees corresponding to windows ≤50 Kb in size and calculated weights using the ‘complete’ method in TWISST (Martin and Van Belleghem 2017). We repeated this analysis on four sets of samples (Table S1): 1) all samples; 2) core range samples (> 380 km from the contact zones); 3) samples from the contact zones or intermediate with core range (i.e., < 300 km from the contact zones); and 4) a set of 14 contact zone samples selected randomly to control for variation because of unequal and small sample sizes. As for ASTRAL, we ran TWISST on autosomal and Z chromosome windows separately.

We incorporated genome-wide recombination rate variation to gain further insights with respect to the processes underlying gene-tree heterogeneity. Regions of low recombination rates should yield more support to the species tree because of reduced local effective population size (*Ne*) in turn reducing the probability of ILS (Pease and Hahn 2013). Further, in low recombining regions barrier loci have a broader effect, resulting in an under-representation of introgressed topologies (Martin et al. 2019). We investigated the relationship between the weight of the 15 possible topologies and local recombination rate. We used estimates of the population recombination rate ⍴ = 4*Ne* × *r* (with *r* the recombination rate and *Ne* the effective population size) in 100 Kb windows (Sebastianelli et al. 2024; available as supplementary data) to attribute a recombination rate to each 500 SNP window. We also used the NJ trees to investigate phylogenetic patterns around the coding sequence of the gene *CYP2J19* (chr8:2377713-2392689), associated with red and yellow forecrown coloration by way of dietary carotenoid conversion (Kirschel et al. 2020). Finally, we inferred genome-wide patterns of introgression by calculating *D* and *Fbranch* statistics based on the autosomal data using Dsuite v0.5r52 (Malinsky et al. 2021) and polarized using the autosomal topology obtained with ASTRAL.

### Analyses of contact zones

To better understand patterns of introgression, we performed geographic cline analyses across the *P. c. chrysconus* v. *P. p. affinis* and *P. c. extoni* v. *P. p. pusillus* contact zones using the ddRADseq datasets. We first investigated changes in genome-wide individual ancestry-proxies along geographic gradients across the contact zones. Lines best representing the center of the contact zones were drawn in ArcGIS v10.7 based on where the two forecrown feather color phenotypes were in approximately equal frequency, and the shortest distance to the line was calculated for each sample. By convention, we considered the contact line as the “0 km” point in the geographic gradient, and negative geographic distances in the RFT side of the contact zone (i.e., either *P. p. affinis* or *P. p. pusillus*). For each sample, we estimated genome-wide ancestry-proxies using ADMIXTURE v1.3 (Alexander et al. 2009) with default parameters and two assumed ancestral parental populations (*K*=2). For this analysis, we used the *populations* module in Stacks to generate vcf files from the outputs of *gstacks*. We filtered SNP loci with vcftools v0.1.16 to keep only biallelic sites with a minor allele frequency ≥0.05, a minimum depth of coverage of 3X, a mean maximum depth of coverage of 14X, and less than 70% missing data (--min-alleles 2 --max-alleles 2 --maf 0.05 --minDP 4 --max-meanDP 14 --max-missing 0.3). We used Hzar v0.2-5 (Derryberry et al. 2014) to fit sigmoid clines to the transition in individual RFT ancestry-proxies along the geographic gradient. We tested four cline models differing by the presence of introgression tails modeling neutral gene flow: 1) tails on either side of the cline, that can be asymmetrical; 2) a single tail on the left side; 3) a single tail on the right side; or 4) no tail at all. As the populations most distant from the contact zones were not admixed (i.e., RFT ancestry=0.0 or 1.0), we did not test models with free maximum and minimum ancestry. We also tested a null model corresponding to no changes in ancestry. We used the corrected Akaike Information Criterion (AICc) to determine the best-fit models. We repeated this analysis for each of the contact zone datasets. We next estimated per-SNP clines based on allele frequencies to test whether the variation in phylogenetic signal across the genome could be linked to variations in the rates of introgression. Based on the output of *gstacks,* we used the *population* module in Stacks to retain only a single SNP per RAD locus (--write-random-snp) and report allele frequencies at each locality (--hzar). Because estimating allele frequencies requires several individuals, localities represented by single samples were lumped with neighboring localities. We subsequently filtered the SNPs to keep only those that: 1) were genotyped in at least two samples in each locality; and 2) were differentially fixed in the two extreme localities of the geographic gradient. We used the resulting SNP sets to model changes in allele frequencies across geographic gradients crossing the contact zones at each locus using the same approach as described above.

### Analyses of mitochondria-interacting nuclear genes

Phylogenomic and cline analyses suggested that the mitochondrial phylogeny (Kirschel et al. 2021), where RFT are monophyletic, best represents the species tree with strong gene flow having eroded the phylogenetic signal in the nuclear genome. We tested whether nuclear-encoded genes interacting with the mitochondria may act as barriers to gene flow and prevent nuclear-mitochondrial genome incompatibilities. We retrieved a list of 208 mitochondrial-interacting nuclear genes from Mikkelsen & Weir (2023), of which we retained 150 that were annotated on the autosomes of the *Pogoniulus pusillus* reference genome (annotation: GCF_015220805.1-RS_2024_03; the list of the 150 genes and their coordinates in the reference genome are given in the Appendix 2). Next, we used the topology weighting results to investigate whether topologies supporting RFT as monophyletic (i.e., as in the mtDNA phylogeny) are more supported in windows overlapping the coding sequences of mitochondria-interacting genes. To determine whether the weight of monophyletic RFT topologies was statistically higher in those regions compared with the genome average, we generated a null distribution of the weights by randomly drawing subsets of 283 windows with replacement, 5,000 times, which corresponded to the number of windows overlapping mitochondria-interacting genes. In a similar way, we tested whether clines in both contact zones were narrower for SNPs located near mitochondria-interacting nuclear genes. Because we did not have a dense coverage of diagnostic SNPs across the genome, we could not recover any SNP located within the coding regions of these genes. Thus, we also considered SNPs located 50 kbp upstream or downstream of the CDS (26 SNPs in Southern Africa, 74 SNPs in Uganda). In both cases, we generated null distributions by randomly drawing 5,000 SNP subsets with replacement.

## Results

### Pervasive gene-tree heterogeneity in the red- and yellow-fronted tinkerbirds radiation

We first inferred evolutionary relationships of the red-and yellow-fronted tinkerbirds using the WGS data. We inferred gene trees in non-overlapping windows and reconstructed the species relationships separately for autosomal and Z chromosome windows while accounting for ILS. The autosomal analysis yielded a fully supported topology (Posterior Probabilities [PP] of all internal nodes=1.0) (Fig. 2a) dividing the red- and yellow-fronted tinkerbirds into four main lineages: *P. c. chrysoconus* (with *P. c. xanthostictus* nested within), *P. c. extoni, P. p. affinis* (with *P. p. uropygialis* nested within), and *P. p. pusillus.* Because species limits do not seem to agree with the traditional two species taxonomy (Kirschel et al. 2021, this study), we hereafter refer to them as *chrysoconus-xanthostictus, extoni, affinis-uropygialis* and *pusillus*, to refer to the northern and southern forms of YFT and RFT respectively. The autosomal topology placed *chrysoconus-xanthostictus* as sister to the other lineages and *affinis-uropygialis* as sister to a monophyletic *pusillus* + *extoni*. Both *uropygialis* and *xanthostictus* were monophyletic within their respective clade and separated by relatively long branches. We repeated this analysis with a subset of 14 samples from core range localities (i.e., > 380 km away from contact zones; hereafter “core range” samples) and found the same relationships (Fig. 2b). Contrastingly, the Z chromosome produced a different topology where *pusillus* was sister to the other lineages and *extoni* sister to a monophyletic *chrysoconus-xanthostictus* + *affinis-uropygialis* (Fig. 2a). The internal branches were very short, suggesting a rapid initial splitting of lineages with low statistical support (PP=0.82) for the sister relationship of *extoni* with *chrysoconus-xanthostictus* + *affinis-uropygialis*. Further, *affinis* and *uropygialis* were reciprocally monophyletic with full support. When analyzing only “core range” samples, we found a different topology where northern taxa *chrysoconus-xanthostictus* and *affinis-uropygialis*, and southern taxa *extoni* and *pusillus,* respectively, were sister taxa (Fig. 2b). All of our reconstructed topologies were discordant with the mtDNA relationships inferred by (Nwankwo et al. 2019).

**Figure 2.**
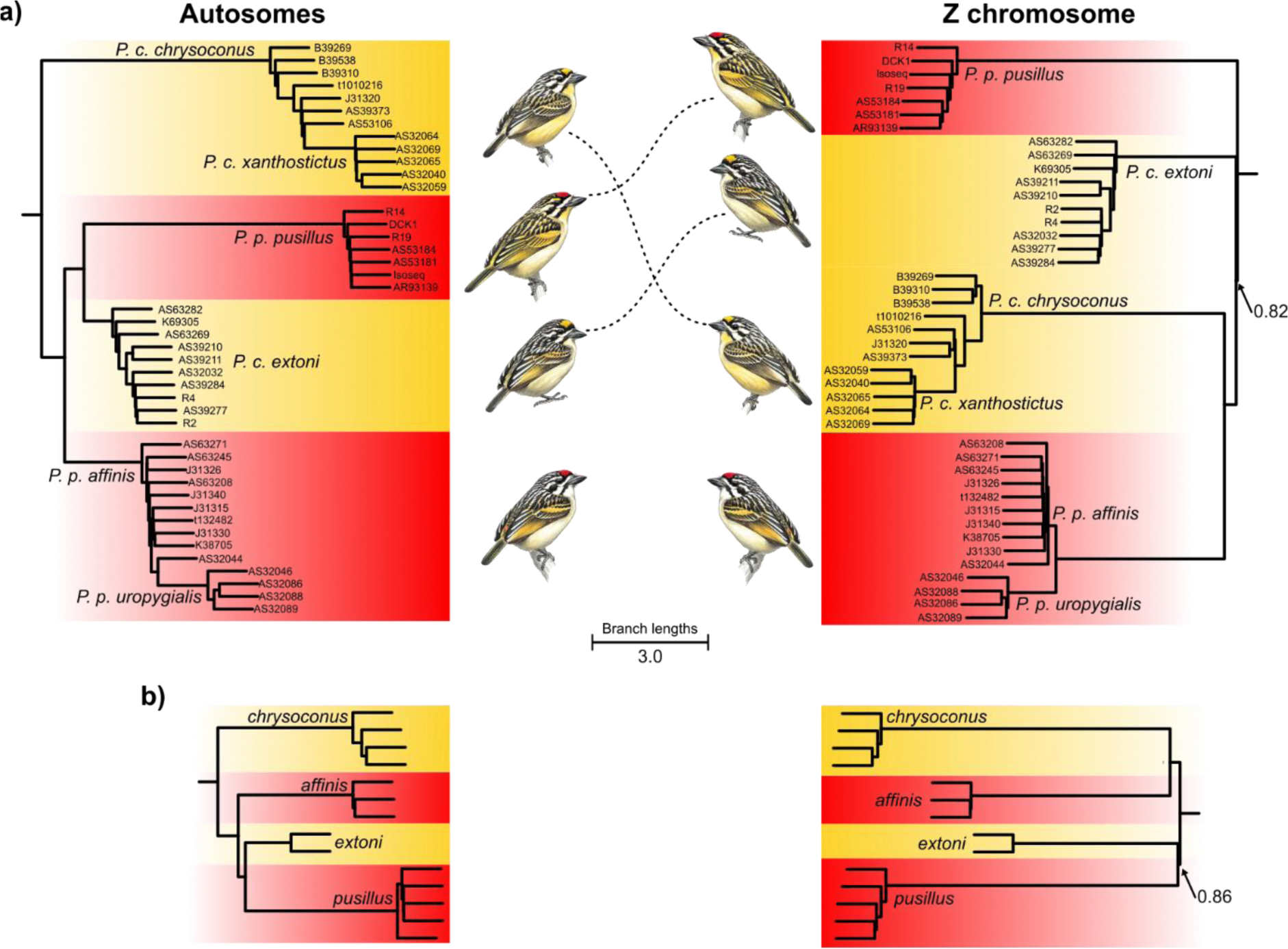
Species trees estimated using ASTRAL. based on Neighbor-Joining trees reconstructed in non-overlapping sliding windows of 500 SNPs using **a)** Autosomal (left) and Z chromosome (right) data from all samples (28,567 and 2871 NJ trees, respectively); **b)** Autosomal (left) and Z chromosome (right) data from “core range” samples only (31,289 and 4154 NJ trees, respectively). All internal nodes received a posterior probability (PP) = 1.0, except for two nodes in the Z chromosome phylogenies, indicated by arrows. Support for nodes within each lineage are not represented for improved clarity. The trees were rooted with sequences of Pogoniulus atroflavus and P. bilineatus, not represented for improved clarity. Branch lengths reflect coalescence units. Tinkerbird paintings courtesy of del Hoyo et al. (2020).

Next, we explored the extent to which gene-tree heterogeneity exists in the red- and yellow-fronted tinkerbird genomes using a topology weighting approach. Based on our genome-wide phylogenetic reconstructions, we merged the *uropygialis* subspecies with *affinis* and separately the *xanthostictus* subspecies with *chrysoconus* to reduce the space of possible species trees. Thus, there are 15 possible rooted species tree topologies, hereafter referred to as topology T1 to T15 (Table S2). We found that the gene trees were overall well resolved with 61.8% of the autosomal windows (n=35,279 out of 57,060) and 90.3% of the Z chromosome windows (n=5,075 out of 5,623) yielding full support (i.e., weight=1, hereafter W1) to a given species tree topology. However, for both chromosomal types, we uncovered important topological heterogeneity with all 15 possible topologies supported by at least one W1 window (Fig. 3a, Table S2). In autosomal windows, five topologies were supported by >8% of the W1 windows in decreasing order of frequency: the topology obtained from the autosome-based phylogenetic inference (T15, 30.1%); (*chrysoconus-xanthostictus*, (*pusillus*, (*affinis-uropygialis*, *extoni*))), (T5, 20.7%); ((*affinis-uropygialis*, *chrysoconus-xanthostictus*), (*extoni*, *pusillus*)), (T3, 13.1%); the Z chromosome topology (T1, 8.4%); and (*chrysoconus-xanthostictus*, (*extoni*, (*pusillus*, *affinis-uropygialis*))) (T8, 8.4%). These topologies all support pairs of monophyletic lineages that also have a secondary contact zone, except for T8 in which RFT are monophyletic (Table S2).

**Figure 3.**
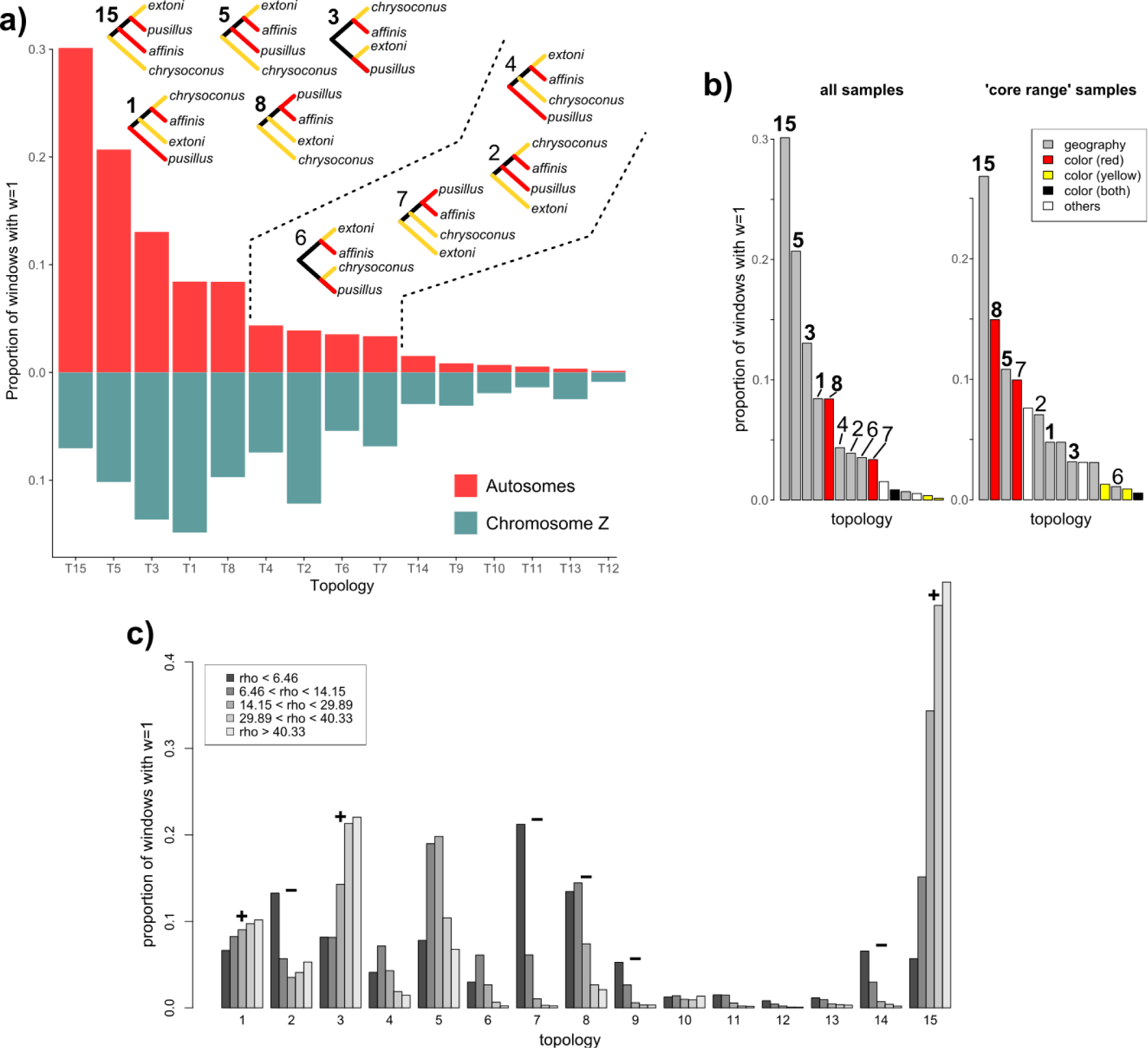
Results of Topology Weighting analyses. a) Proportion of windows with a weight=1 supporting each of the 15 possible species trees in the autosomes (red), and chromosome Z (green). The topologies are ordered based on their support in the autosomal analysis, and the nine with the highest proportion of windows are illustrated. b) Autosomal results based on two sample subsets: all samples (left barplot, same as red bars in a) and 14 “core range” samples collected far from the contact zones. The topologies are colored according to one of five categories defined based on the taxa supported as monophyletic (see Table S2 for more details). c) Variation of the proportion of autosomal windows with weight=1 at each topology depending on the recombination rate (represented here as the population recombination rate, rho). Windows were divided into five categories corresponding to the 0-5%, 5-20%, 20-80%, 80-95% and 95-100% quantiles of the recombination rate distribution. Symbols above the bars denote topologies that either increase (+) or decrease (-) in support with increasing recombination rate.

Plumage colouration appeared to be a poor predictor of phylogenetic relationships. The topologies that supported RFT as monophyletic had relatively low frequencies (T7, T8, T9, accounting for a total of 12.6% of W1 windows) and YFT was rarely monophyletic (T9, T12, T13, total of 1.4% of W1 windows). Topological variation on the Z chromosome was very distinct from that of autosomes (Fig. 3a). There, the most commonly supported topologies were T1 (14.9%), T3 (13.7%), and (*extoni*, (*pusillus*, (*affinis-uropygialis*, *chrysoconus-xanthostictus*))) (T2, 12.2%). All three of these topologies supported *affinis-uropygialis* as sister species to *chrysoconus-xanthostictus*. Topologies T5 (10.2%) and T8 (9.7%) were also supported by >8% of W1 windows on the Z chromosome. The most striking difference between the autosomes and the Z chromosome was the support for T15, which dropped from 30.1% in the former to 7% in the latter. As in the autosomes, the topologies where RFT was monophyletic were more frequent than those where YFT was monophyletic (19.7% vs 6.4% of the W1 windows, respectively).

In the autosomes, we did not find large contiguous blocks of windows supporting the same topology. Instead, these windows were dispersed along the chromosomes, interspersed with windows supporting other topologies (Figs. 4a, S1). Contrastingly, on the Z chromosome, we identified four contiguous blocks of windows spanning 2.7-6.4 Mb with a high support for T1 & T3. The cumulative support for monophyletic RFT topologies was also higher on the Z chromosome compared with the autosomes, with a 3.5 Mb block supporting these topologies almost exclusively.

**Figure 4.**
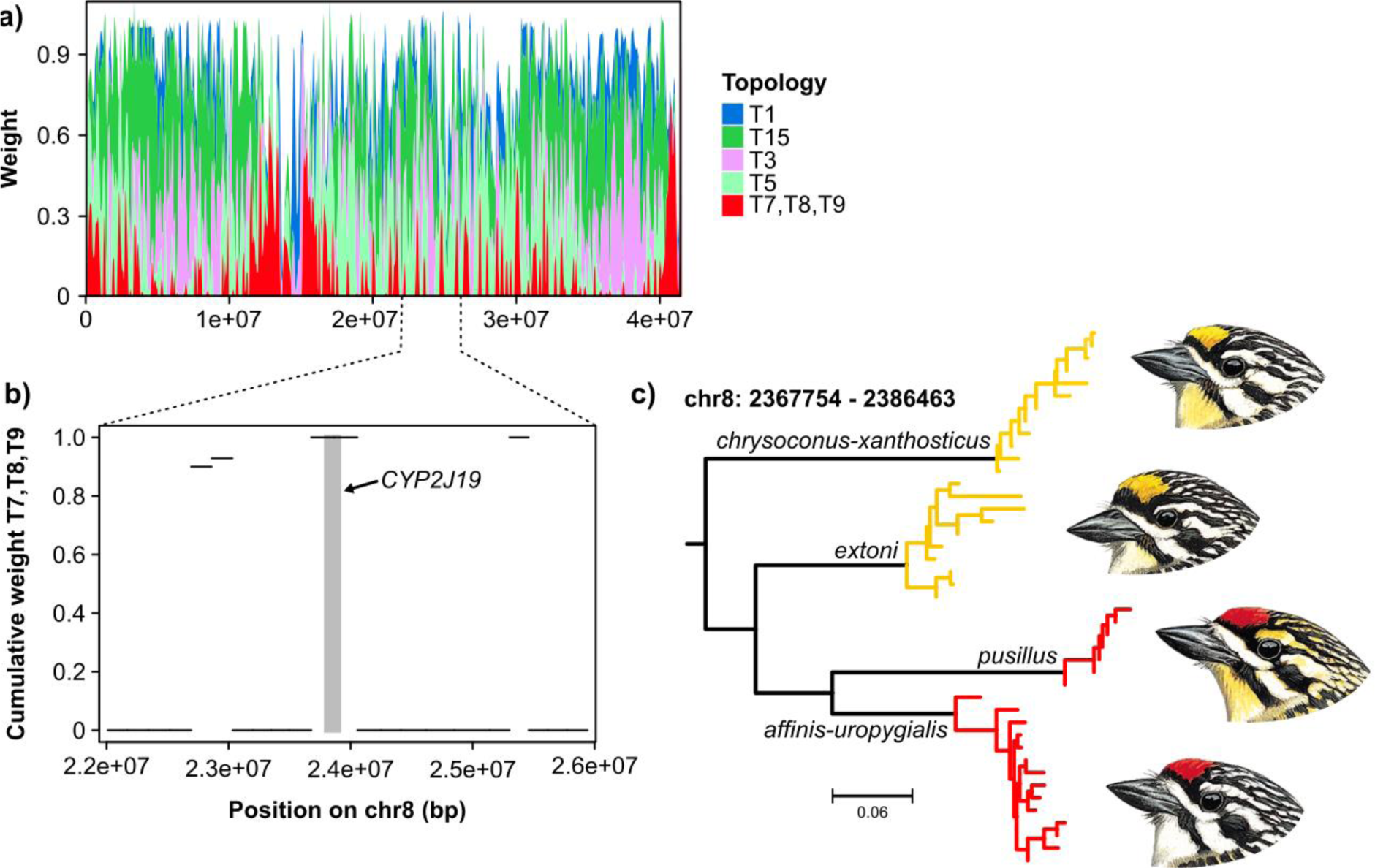
Phylogenetic signal on chromosome 8. a) Distribution of the weights of the four commonest topologies (T1, T3, T5 and T15) as well as cumulative weights of the topologies where Red-fronted Tinkerbirds are monophyletic (T7, T8 and T9) along chromosome 8. The weights were smoothed using a local regression with a span of 0.01. b) Zoom into the region surrounding the gene CYP2J19 (gray rectangle shows the coding sequence of the gene), with the horizontal bars representing the cumulative weight of T7, T8 and T9 in each 500 SNP window. c) Neighbor-joining phylogenetic tree of the first window overlapping the coding sequence of CYP2J19. Tinkerbird paintings courtesy of del Hoyo et al. (2020).

We classified if the topologies were concordant according to 1) forecrown plumage coloration, or 2) whether sister lineages reflected their geographical distribution (i.e., lineages with a contact zone were sister; hereafter “geography”) (Table S2). The “geography” category dominated the signal with 81.3% and 67.3% of the W1 autosomal and chromosome Z windows, respectively (Figs. 3b, S2, S3). To determine whether these levels of heterogeneity could be recovered even when considering only samples from localities far away from the contact zones, we partitioned the dataset into two subsets, “core range” (14 samples) and “contact zones” (29 samples), and repeated the analysis. Considering the imbalance in sample size where most samples were located proximal to the contact zones, we also generated a “control” subset of 14 samples from the contact zones to match the sample size of the “core range” subset. The “contact zones” subset produced nearly identical results as the complete set; however, two monophyletic RFT topologies (T7 & T8) in the “core range” subset increased substantially in frequency in both autosomal and Z chromosome analyses (Figs. 3b, S3). The “contact zones” control subset showed that with a reduced sample size, even when using samples from the contact zones, T7 and T8 were more frequent than when using all the samples (Figs. S2, S3). Importantly, the mean weight of T7 and T8 was significantly higher when using the “core range” samples compared with the control subset (Wilcoxon rank sum test *p*<2.2×10^−16^ in both cases). Despite these differences, T15 was the dominant topology in the autosomes and T1 & T3 in the Z chromosome data, with considerable gene-tree heterogeneity in all analyses.

We quantified excesses in derived allele sharing across lineage pairs to determine whether gene-tree heterogeneity could be attributed to introgression. We calculated *D* statistics and *f4* ratios for all four trios using the same aforementioned lineages. When arranging the trios according to the ASTRAL topology, we found a significant excess of derived allele sharing in all four of them (Table 1). The strongest signal of introgression was recovered in trios with either *extoni* or *pusillus* as P1, *affinis-uropygialis* as P2 and *chrysoconus-xanthostictus* as P3 (*D*=0.22 and 0.18, respectively). The two other trios supported weak introgression between *affinis-uropygialis* and *extoni*, and *pusillus* and *chrysoconus-xanthostictus,* respectively (*D*=0.03 in both cases). We next calculated *Fbranch* statistics from the *f4* ratios; however, this analysis did not recover introgression involving internal branches of the tree (Fig. S4).

**Table 1.** D-statistics calculated from autosomal SNPs based on the species tree topology inferred with ASTRAL.

| P1 | P2 | P3 | D | Z-score | N<sub>BBAA</sub> | N<sub>ABBA</sub> | N<sub>BABA</sub> |
| --- | --- | --- | --- | --- | --- | --- | --- |
| *extoni* | *affinis* | *chrysoconus* | 0.22 | 95.65 | 912,934 | 716,947 | 458,258 |
| *pusillus* | *affinis* | *chrysoconus* | 0.18 | 80.70 | 871,476 | 714,146 | 490,317 |
| *pusillus* | *extoni* | *affinis* | 0.03 | 9.43 | 708,725 | 730,86 | 691,286 |
| *extoni* | *pusillus* | *chrysoconus* | 0.03 | 14.62 | 957,302 | 558,730 | 523,679 |

### Monophyly among parapatric lineages overrepresented in high recombination rate regions

When considering only regions with very low recombination rate (population recombination rate (⍴) < 6.46, corresponding to the 5% quantile of ⍴), the monophyletic RFT topologies T7 and T8 were the most common (23% and 13% of the W1 windows, respectively). With increasing recombination rate, their proportion rapidly decreased while some topologies that included geographically proximate populations increased (e.g., T1, T5 and T15). For four topologies, support decreased with increasing recombination rate: T7, T8, T9 and T14. T2 also decreased in the first three categories, but slightly increased in the highest recombination rate categories. Topologies T7, T8 and T9 supported RFT as monophyletic, and topologies T2, T7 and T14 supported *extoni* as sister to the remaining taxa. On the contrary, three topologies showed a strong increase in frequency: T1, T3 and T15. These all supported *pusillus* + *extoni*, *affinis-uropygialis* + *chrysoconus-xanthostictus*, or both, as monophyletic. We subsequently used windows with the lowest recombination rate for phylogenetic reconstruction and found that topology T7 was fully supported (Fig. S5), which is concordant with mitochondrial relationships (Nwankwo et al. 2019; Kirschel et al. 2021) and suggests a monophyletic RFT.

We explored the phylogenetic signal around the gene *CYP2J19*, underlying red and yellow colorations (Kirschel et al. 2020), to investigate whether it supported a single origin of red forecrown, as suggested by our preferred species tree hypothesis. We found that the two windows overlapping the coding sequence of *CYP2J19* (chr8: 2367754-2386463 and chr8: 2386466-2405472) fully supported topology T8 where RFT is monophyletic (Figs. 4b, 4c). Contrastingly, 23 out of 26 windows flanking the gene (i.e., 250kbp downstream or upstream the coding region) supported paraphyly for RFT (Fig. 4b). Only two windows upstream and one downstream carried a weight >0.8 for T7, T8 and T9 considered together.

### Levels of introgression across contact zones are associated with phylogenetic signal

Our analyses suggest that RFT taxa *pusillus* and *affinis-uropygialis* are sister lineages, and that introgression between parapatric lineages generated high levels of phylogenetic heterogeneity. It remains unclear, however, whether these patterns of introgression are the legacy of ancestral (i.e., at the first stage of secondary contact) or contemporary gene flow. To clarify this, we used the ddRADseq dataset to study transitions in genomic ancestry and allele frequencies along geographic gradients that cross two contact zones: between *chrysoconus* and *affinis* (in Uganda and Kenya), and between *extoni* and *pusillus* (in Southern Africa). Clines fitted to ancestry-proxies (i.e., *Q* values) were narrow (Figs. 5a, 5d; cline parameters are detailed in Table S3), with widths of 10 km (95% CI [6.8, 15.6]) for the *chrysoconus* v. *affinis* cline in the north and 4.3 km (95% CI [1.2, 14.2]) for the *extoni* v. *pusillus* cline in the south. However, we inferred bi-directional introgression tails in the latter, which suggested neutral introgression, with admixed individuals found >30km from the contact zone. By contrast, the northern contact zone was best modeled by a simple cline without introgression tails.

**Figure 5.**
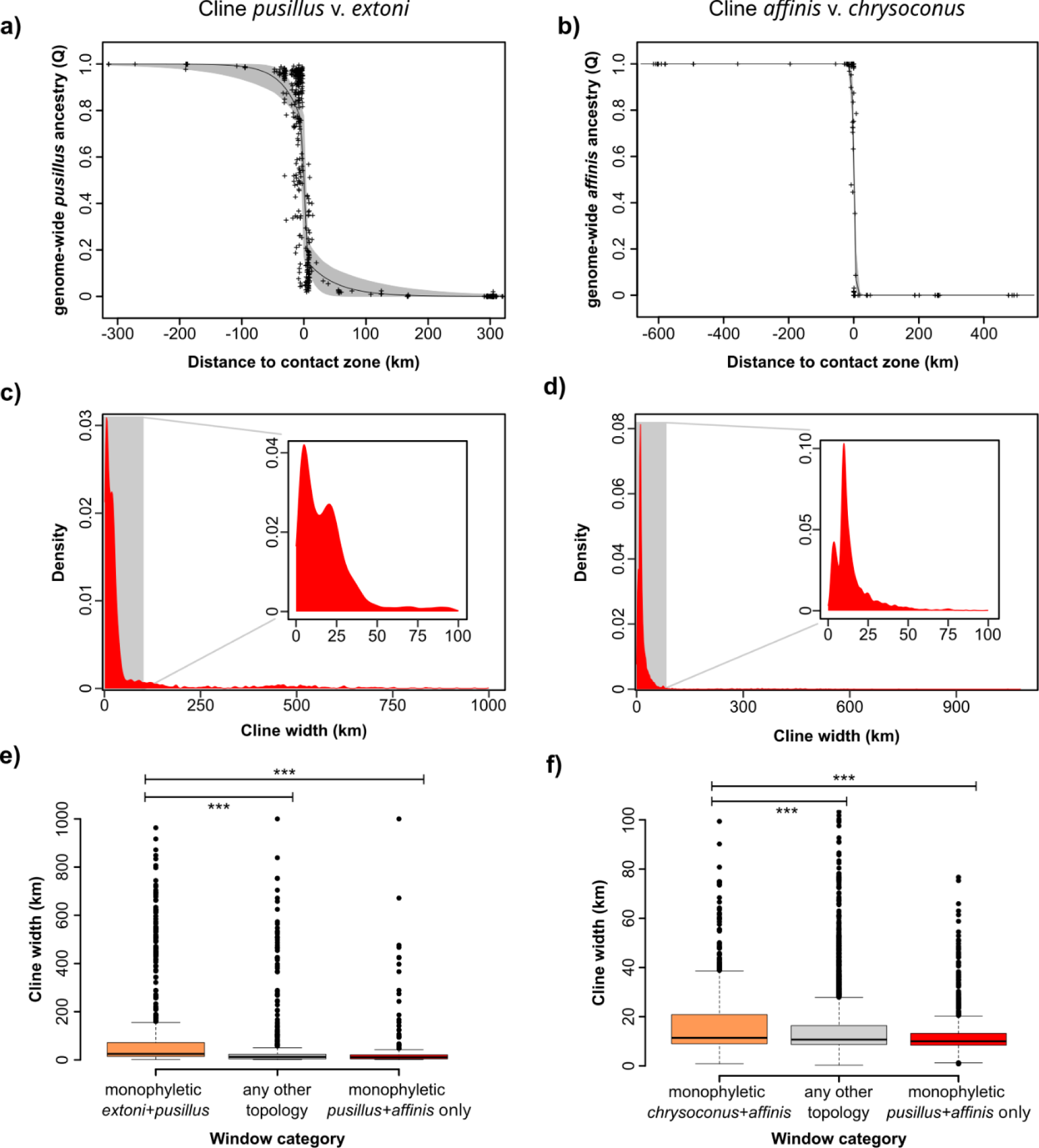
Results of cline analyses across the extoni v. pusillus (a,c,e) and the chrysoconus v. affinis (b,d,f) contact zones. a-b) Sigmoid clines fitted to genome-wide individual ancestry-proxies (Q values) calculated using ADMIXTURE, represented as black crosses. Black lines show the maximum likelihood cline, and the gray envelope the 95% credibility parameter space. c-d) Density plots of the distribution of cline widths in single locus analyses. Insets show density distributions for loci with a cline width ≤100 km. e-f) Distribution of cline widths in genomic regions supporting three categories of topologies (from left to right): topologies supporting species with a contact zones (i.e., either chrysoconus-xanthostictus + affinis-uropygialis or extoni + pusillus) as monophyletic; topologies supporting any other taxa pair as monophyletic (“all others”, includes 12 topologies); of the latter, those specifically supporting pusillus + affinis-uropygialis as monophyletic (T7, T8 and T9). Note: in f), the y-axis range was shortened to 100 km to improve visualization, but the statistical analysis was run on the complete data range.

We next fitted sigmoid clines to the transitions in allele frequencies at diagnostic SNPs (i.e., SNPs that are differentially fixed in the populations most distant from the contact zone) and tested whether some topologies were associated with wider clines than others. If patterns of gene tree heterogeneity are the result of contemporary gene flow, we predict that genomic regions supporting species with a contact zone as monophyletic would also harbor SNPs with wider clines. Conversely, clines should be narrower in regions supporting the putative species tree topology (with *affinis-uropygialis* and *pusillus* as sister lineages). In Southern Africa (*extoni* v. *pusillus* transition), the mean width of single-locus clines was 65.23 ±SD 144.09 km (range=1.3-999.8 km) (Fig. 5c). We compared cline widths at SNPs located in genomic regions that supported three categories of topologies: *extoni*+*pusillus* as sister taxa (topologies T3, T10 and T15); any other taxa pair as sister taxa (i.e., all other topologies except the aforementioned three); and specifically *affinis-uropygialis* and *pusillus* as sister taxa (topologies T7, T8 and T9, also included in the previous category). We found that SNPs in the *extoni*+*pusillus* category followed significantly wider clines (mean width=102.76 km) than those in the other two categories (width=39.27 and 33.93 km, respectively; Wilcoxon rank sum test *p*<2.2×10^−16^), even when we removed outlier loci (i.e., clines with a center>100 km away from the contact zone or a width>300 km) (Figs. 5e, S6). Across the Uganda / Kenya contact zone (*chrysoconus* v. *affinis* transition), single-locus clines were significantly narrower (mean 25.39 ±SD 67.43 km, range=0.3-1080.8 km) than in Southern Africa (Wilcoxon rank sum test p<2.2×10^−16^) (Fig. 5d). Again, we divided the SNPs in three categories depending on local phylogenetic support (*chrysoconus*+*affinis*: topologies T1, T2 and T3; all other topologies; *affinis*+*pusillus* only). SNPs in the *chrysoconus*+*affinis* category followed significantly wider clines than those in the “all other topologies” category (mean widths=17.96 and 15.72 km, Wilcoxon rank sum test *p*=2.8×10^−4^) (Figs. 5f, S7). The difference between the *chrysoconus*+*affinis* and *affinis*+*pusillus* categories was more marked (width=12.00 km, *p*=5.45×10^−13^). Finally, we found a positive correlation between cline width and recombination rate in both contact zones (Spearman rank correlation test *p*<2.2×10^−16^, Rho=0.28 and 0.47) (Fig. S8).

### Mitochondria-interacting genes are not barriers to gene flow

Incompatibilities between the mitochondrial and nuclear genomes have been proposed as an important driver of reproductive isolation (Hill 2016). Thus, if the mitochondrion accurately represents the tinkerbird species tree (i.e., monophyly of RFT) and the nuclear genome is largely introgressed, we hypothesized that mitochondria-interacting nuclear genes (N-Mt genes) would act as barriers to gene flow to avoid detrimental combinations of mitochondrial and nuclear genes. To test this hypothesis, we investigated the phylogenetic signal in 283 500-SNP windows that overlapped the coding regions of 150 N-Mt genes. The topological support in these regions was overall similar to genome-wide levels, with T15, T5 and T3 the most common topologies (Fig. S9a). The weight of monophyletic RFT topologies (mean weight=0.13) in windows that overlapped N-Mt genes was not significantly higher than in 5,000 random draws with replacement of 283 windows (mean weight=0.12, 95% quantile=0.15) (Fig. S9b). None of the SNPs included in the cline analysis were located within coding regions of N-Mt genes. However, we also considered 50 Kb of sequence that flanked the targeted N-Mt genes. We recovered 38 SNPs in the *extoni* / *pusillus* dataset and 74 SNPs in the *chrysoconus* / *affinis* dataset. These SNPs did not support narrower clines than random subsets of the same size (southern Africa: mean width=70.65 km, compared with a background mean=75.88 km with 5% quantile=31.91 km; Uganda / Kenya: mean width=36.59 km compared with a background mean=27.51 km with 5% quantile=15.14 km) (Fig. S10).

## Discussion

Gene flow and admixture have a crucial role in evolution, both as forces counter-acting local adaptation and divergence (Kawecki and Ebert 2004), and as sources of novel allele combinations fueling diversification (Meier et al. 2023). The temporal stage at which such genetic exchange occurred is critical to understanding how barriers to gene flow evolve and shape genome-wide divergence. In this study, we investigated introgression in red- and yellow-fronted tinkerbirds (*Pogoniulus pusillus-chrysoconus* species group), a complex of closely related barbets with intricate secondary contact zones across Africa (Nwankwo et al. 2019; Kirschel et al. 2020; Kirschel and Sebastianelli 2022; Sebastianelli et al. 2024), by characterizing patterns of phylogenomic variation. Using genome-wide SNPs, we recovered a fully supported topology where neither currently recognized species was monophyletic. Instead, the yellow-fronted *P. c. chrysoconus* diverged earliest in the radiation, followed by the red-fronted *P. p. affinis,* which was sister to two forms from Southern Africa (red-fronted *P. p. pusillus* and yellow-fronted *P. c. extoni*) that are geographically adjacent to one another. The subspecies *P. c. xanthostictus* and *P. p. uropygialis* were nested within *P. c. chrysoconus* and *P. p. affinis,* respectively. Although this topology was fully supported, the phylogenetic relationships of these four lineages varied across the genome. Topologies placing taxa with a secondary contact zone as sister to each other (e.g., *chrysoconus*+*affinis, extoni*+*affinis*, *extoni*+*pusillus*) were supported by 80% of the genomic windows. The only exception was the pair of yellow-fronted *chrysoconus* + *extoni*, which presumably have been in recent contact in Uganda (Kirschel and Sebastianelli 2022) yet were rarely monophyletic (1.4% of the windows). By contrast, RFT were more frequently recovered as monophyletic. This discord between phenotypic similarity and genome-wide phylogenetic signal suggests either parallel evolution of red and yellow forecrown coloration, or a genome-wide erosion of the phylogenetic support for the species tree, i.e. due to introgression.

### Neutral gene flow through narrow hybrid zones shapes gene-tree heterogeneity

The extent of phylogenetic heterogeneity across the genome as a consequence of introgression depends on the biogeographic history of the populations, as well as processes acting on the genome, in particular natural selection and genetic recombination (Burbrink and Gehara 2018; Musher et al. 2024; Thom et al. 2024). Theory anticipates a positive correlation between introgression and recombination rate is produced when introgressed alleles are purged by selection (Schumer et al. 2018; Martin et al. 2019; Duranton and Pool 2022). Such a correlation has been used to identify local phylogenetic signals that likely result from introgression (Li et al. 2019; Hennelly et al. 2021; Foley et al. 2024; Jiang et al. 2024). We found that the phylogenetic relationships across RFT and YFT were strongly associated with recombination rate, where genomic windows of lower recombination rates supported the monophyly of RFT. By contrast, two of the dominant topologies in the genome-wide analyses (T3 & T15) were relatively rare in windows characterized by low recombination rates, and their frequencies were strongly positively correlated with recombination rate. It should be noted that a negative relationship between introgression and recombination rate could potentially arise, particularly when the introgressed alleles are under positive selection (Duranton and Pool 2022; Feng et al. 2024). However, our data showed a positive correlation, supporting that introgressed variants are mostly neutral or selected against. Regions of low recombination supported a topology where *P. c. extoni* splits first, and *chrysoconus*-*xanthostictus* are sister to a monophyletic RFT. We believe this best represents the red- and yellow-fronted tinkerbirds species tree and introgression between parapatric lineages drives gene-tree heterogeneity, even in strictly allopatric populations.

We modeled changes in genome-wide ancestry-proxies (*Q* values) for two of the contact zones and estimated cline widths of 4-10 km, with such narrow transition zones comparable to other avian systems where strong reproductive isolation has been inferred (Price 2008; Grossen et al. 2016; Pulido-Santacruz et al. 2018). As cline width depends on dispersal rates and selection against hybrids, a narrow cline reflects strong reproductive isolation (Barton and Hewitt 1985; Payseur 2010; Dufresnes et al. 2021). Despite that, we detected introgressed alleles hundreds of kilometers away from both contact zones. As reproductive barriers build up across a genome, the distribution of cline width at independent loci is expected to change from a normal distribution to a Poisson distribution, and finally a bimodal distribution where many loci are impermeable to gene flow (Dufresnes et al. 2021). Here, we report distributions of cline widths similar to Poisson distributions, with the emergence of two modes at lower widths. Furthermore, a model with introgression tails best represented the *extoni* v. *pusillus* transition, consistent with neutral introgression far from the contact zone (Szymura and Barton 1986). Thus, despite narrow transitions in ancestry-proxies, tinkerbird hybrid zones appear porous to neutral introgression, a conclusion that is further supported by the positive relationship between cline width and recombination rate (Duranton and Pool 2022). Introgression dynamics differed across the two contact zones analyzed. The absence of introgression tails in the *chrysoconus* v. *affinis* genome-wide analysis and relatively narrower single SNP cline widths suggests an overall lower rate of introgression in their contact zone, consistent with the phylogenetic analyses that support *extoni* and *pusillus* as sister lineages.

### Phylogenetic signal for the species tree is conserved in small, interspersed genomic regions

In both contact zones, the genomic regions that supported parapatric taxa as monophyletic also produced wider transitions in allele frequencies. Because we considered markers that are differentially fixed in allopatric populations on both sides of the contact zone, intermediate allele frequencies along the transect are maintained by ongoing gene flow rather than ancient admixture. Further support is that topology T15 remains dominant even when only allopatric samples were considered, with the *extoni* v. *pusillus* ancestry cline revealing broad introgression tails. Thus, our phylogenomic and geographic cline analyses draw a coherent picture where the dominant phylogenetic signal results from contemporary introgression across contact zones, while support for the species tree is conserved in small and interspersed genomic regions. In the *chrysoconus* v. *affinis* contact zone, two topologies that do not support *chrysoconus-xanthostictus* + *affinis-uropygialis* as monophyletic (T5 and T15) were associated with wide clines; however, these topologies are over-represented in regions of medium or high recombination rate, respectively, which are more permeable to introgression in general. Considering that there are several parallel RFT and YFT hybrid zones across Africa, phylogenetic signal at a given locus must depend on the relative strength of introgression in the different hybrid zones. In this specific case, T5 supports *extoni* + *affinis-uropygialis* as monophyletic, and T15 *extoni* + *pusillus.* While we did not focus specifically here on the *extoni* v. *affinis* transition, we found that introgression between *extoni* and *pusillus* was stronger than between *chrysoconus* and *affinis*, consistent with the hypothesis that T15 overrides phylogenetic signals in many regions with high recombination rate. Finally, one limitation of our approach is that we could not take recombination breakpoints into account when inferring trees, thus these local phylogenies may not accurately represent the evolutionary relationships of haplotypes (Martin and Van Belleghem 2017). Consequently, a given local phylogeny likely reflects the dominant signal of introgression in the corresponding genomic window if it overlaps several haplotypes. This may also result in an under-estimation of ancient introgression compared with contemporary gene flow as recombination gradually breaks down introgressed haplotypes (Racimo et al. 2015).

Introgression across species is commonly reported in phylogenomic studies (Mallet et al. 2016; Hibbins and Hahn 2022), including cases where the dominant phylogenetic signal results from introgression (Fontaine et al. 2015; Zhang et al. 2021). Here, we show that neutral introgression through narrow secondary contact zones contributes to gene-tree heterogeneity, to a point that introgressed topologies are vastly dominant, even in the species’ core ranges. This is consistent with a study on the suboscine bird radiation (Singhal et al. 2021), where range overlap was among the main predictors of the levels of introgression. However, there are examples of closely related, parapatric lineages that do not show introgression (Pulido-Santacruz et al. 2020; Sarver et al. 2021; Khan et al. 2024), while deeply diverged parapatric lizard species do (Caeiro-Dias et al. 2021). In that context, the nature of reproductive barriers may be the most important factor constraining introgression, and thus shaping the genomic landscape of gene-tree heterogeneity. Because of that, introgression must be carefully taken into account when reconstructing the phylogenetic relationships of species with range overlap, particularly bearing in mind that the most common topology genome-wide may not represent the species tree. While the identification of the species tree in these conditions remains challenging, systematic investigations of genome-wide phylogenetic variation in the light of recombination rate may yield more accurate phylogenetic reconstructions and a better understanding of the factors underlying gene-tree heterogeneity.

### Red- and yellow-fronted tinkerbirds maintain phenotypic integrity despite gene flow

One important implication of our results is that RFT and YFT lineages can withstand very high levels of introgression across their whole ranges while maintaining their phenotypic integrity and genome-wide differentiation. Mito-nuclear incompatibilities have been proposed as an important driver of reproductive isolation (Hill 2016; Burton 2022), because co-adaptation of mitochondrial and mitochondria-interacting nuclear genes may cause hybrid disfunction (Chou and Leu 2010). While mitochondrial introgression is very common in animals (Toews and Brelsford 2012), we found that the mitochondrial relationships of the red- and yellow-fronted tinkerbirds likely reflect the species tree rather than introgression. It would be tempting to interpret this as evidence that mito-nuclear incompatibilities across species prevent mitochondrial introgression. However, we did not find evidence of reduced introgression around the coding sequences of 150 mitochondria-interacting nuclear genes, which contradicts this hypothesis. Similarly, (Mikkelsen and Weir 2023) did not find evidence that mitochondria-interacting nuclear genes co-introgressed along the mitochondrial genome in recently diverged skua species. This may mean that, in the cases of both tinkerbirds and skuas, allopatric divergence did not last long enough for mitonuclear incompatibilities to build up (Mikkelsen and Weir 2023), although it has been argued that they could arise at short time scales (Barnard-Kubow et al. 2016; Hill 2016). Alternatively, mitonuclear incompatibilities may not play an important role in reproductive isolation, consistent with the sparsity of empirical evidence supporting this phenomenon (Burton 2022). There are of course scenarios where the mitochondrial genome might not introgress across a species boundary because of processes unrelated to mito-nuclear incompatibilities, e.g. because of male-mediated gene flow (Peters et al. 2012; Toews and Brelsford 2012) or female-biased host specificity (Spottiswoode et al. 2011).

Previous investigations of *extoni* and *pusillus* in Southern Africa suggest that variation in forecrown color and song rhythm might instigate assortative mating (Kirschel et al. 2020; Sebastianelli et al. 2024). There, the phenotypic and genome-wide transitions are similarly sharp (Nwankwo et al. 2019; Sebastianelli et al. 2024), demonstrating that introgression does not compromise the phenotypic integrity of *pusillus* and *extoni*. The levels of gene-tree heterogeneity that we report across the whole species complex suggest that similar mechanisms play out in the other contact zones. The genetic architecture of forecrown color and song rhythm appears relatively simple, with respectively one and four genes specifically identified so far (Kirschel et al. 2020; Sebastianelli et al. 2024). Thus, it is possible that reduced introgression at these loci combined with assortative mating contribute to maintaining phenotypic integrity in allopatric populations, while the genomic background is largely introgressed. Under this hypothesis, recombination would break the association between alleles associated with phenotypic traits and introgressing neutral variants (Barton and Bengtsson 1986; Martin and Jiggins 2017; Schumer et al. 2018), which is consistent with the genomic distribution of topology weights in our analysis, where many small windows that support RFT monophyly are interspersed across the genome. Particularly, two such windows overlap the gene *CYP2J19* on Chromosome 8, which has been identified as the ketolase that catalyzes the conversion of dietary yellow carotenoids to red ketocarotenoids and thus explains differences in forecrown coloration (Kirschel et al. 2020). This finding lends further support to the hypothesis that genes underlying phenotypic traits involved in reproductive isolation resist introgression, and are decoupled from introgressed variants by recombination. The local phylogenies around *CYP2J19* further support the RFT as monophyletic, along with a single origin of the red forecrown coloration, contrastingly with the genome-wide phylogenetic signal. Further investigations, especially phenotype-genotype association analyses, will clarify whether the same mechanisms underlie the red forecrown color in the two RFT lineages.

Although our results shine a new light on speciation mechanisms and the evolutionary history of red- and yellow-fronted tinkerbirds, several phenomena remain unexplained. Our topology weighting analyses revealed contrasting patterns in the autosomes and the Z chromosome. In the latter, the phylogenetic signal for a monophyletic *extoni* + *pusillus* was much reduced, while several large clusters of windows supported monophyly in either the northern taxa *chrysoconus-xanthostictus* + *affinis-uropygialis,* or the red-fronted taxa *affinis-uropygialis + pusillus.* This pattern suggests differential introgression on the Z chromosome across the different hybrid zones. Sex chromosomes play a major role in speciation, because their reduced recombination facilitates the emergence of reproductive incompatibilities (Irwin 2018). Because of that, sex chromosomes are generally considered to be less permeable to gene flow compared to autosomes (Muirhead and Presgraves 2016; Fraïsse and Sachdeva 2021), and have been used to identify the correct species relationships in the presence of introgression (Fontaine et al. 2015; Johnson et al. 2023; Jiang et al. 2024). However, cases of sex chromosome introgression have also been reported (Li et al. 2016; Jensen et al. 2024; Rancilhac et al. 2024). Further investigations will be required to clarify the role of sex chromosomes in tinkerbird speciation. Another puzzling aspect is the maintenance of ecological differences in the face of gene flow, as the four RFT and YFT lineages occupy distinct habitats and occur along ecological gradients (Kirschel et al. 2021; Sebastianelli et al. 2024). In this context, introgression can be deleterious by acting against local adaptation (Kawecki and Ebert 2004). Similarly to phenotypic traits, the genetic architecture of local adaptation likely plays a key role in its maintenance in the face of gene flow (Tigano and Friesen 2016).

### Conclusions

In this study, we combined phylogenomics and hybrid zone analyses in a group of closely related, parapatric bird species to understand the contribution of contemporary gene flow to gene-tree heterogeneity, and its relevance to understanding speciation mechanisms. Our results highlight that the genomes of red- and yellow-fronted tinkerbirds are mosaics of regions supporting different topologies as a result of introgression. Particularly, we demonstrate that contemporary gene flow through hybrid zones contributes to this pattern. Using genomic regions with the lowest recombination rate appears to be a relevant approach to identify the species relationships in the face of introgression.

Red- and yellow-fronted tinkerbirds appear to represent one more example where introgression results in the dominant phylogenetic signal being discordant with the species tree. This conclusion is especially important when investigating the evolution of forecrown coloration: while the genome-wide topology suggests parallel evolution of red and yellow forecrowns, our favored phylogenetic scenario based on investigation of gene-tree heterogeneity supports a single origin of red feather color. Thus, our study further demonstrates the relevance of phylogenetic analyses of whole genome data to understand the evolution of phenotypic traits, and consequently, reproductive isolation.

## Acknowledgements

We thank L Hadjioannou, EC Nwankwo, M Mamba, TG Hadjikyriakou and KG Mortega for fieldwork, A Howland, S Agostini, G Vogt, the landowners of Kube Yini Private Game Reserve and of Seringveld, De Tweedespruit and Cullinan Conservancies (especially E and J Fourie, J Visser, L Klapwijk, J Du Toit and R Olivier), S Willows-Munro, C Niesler, N and L Baker, R Mulwa, D Pomeroy and A Monadjem for logistical support, and Louisiana State University Museum of Natural Science (DL Dittmann) the Zoological Museum of the University of Copenhagen (J Bolding Kristensen), and the Center for Tropical Research at the University of California Los Angeles (TB Smith) for providing samples. The Ethiopian Wildlife Conservation Authority, Uganda Wildlife Authority, Uganda National Council for Science and Technology, Kenya Wildlife Service, Kenya’s National Council for Science and Technology, National Museums of Kenya, Tanzania Wildlife Research Institute (TAWIRI), Tanzania Commission for Science and Technology (COSTECH), Tanzania National Parks (TANAPA), Eswatini Big Game Parks, Ezemvelo KZN Wildlife, Mpumalanga Tourism and Parks Agency, Gauteng Department of Agriculture and Rural Development (GDARD), and the Animal Research Ethics Committee of the University of KwaZulu-Natal (ref: AREC/00002381-82/2021), and the Cyprus Ministry of Agriculture, rural development and environment veterinary services provided research permits. Funding was provided by the European Regional Development Fund and the Republic of Cyprus through the Research and Innovation Foundation (Projects: EXCELLENCE/0421/0413 and EXCELLENCE/0421/0301), University of Cyprus Research Grant No, 8037P-25023, and FP7 Marie Curie Reintegration Grant No. 268316 to A.N.G.K., and A.G. Leventis Foundation scholarships to S.G.d.S., S.M.L, M.S. and M.M.

## Author contributions

**L.R.**: Conceptualization, Formal Analysis, Methodology, Visualization, Writing - Original Draft Preparation, Writing - Review & Editing; **S.G.dS**., **B.O.O.**: Data Curation, Writing - Review & Editing; **S.M.L.**, **M.S., T.A., C.T.D.**: Investigation, Resources, Writing - Review & Editing; **M.M.**, **C.N.**: Investigation, Writing - Review & Editing; **A.B.**: Methodology, Validation, Writing - Review & Editing; **B.M.vH.**: Conceptualization, Project Administration, Resources, Methodology, Validation, Writing - Review & Editing; **A.N.G.K.**: Conceptualization, Investigation, Funding Acquisition, Project Administration, Resources, Writing - Review & Editing.

## Data availability statement

The supplementary tables and figures, Appendix 1 & 2 and the data files necessary to reproduce the analyses and figures are available at doi:10.5061/dryad.0cfxpnwb3 (https://datadryad.org/stash/share/fNwX3znR08TK7henNPNLwPFXfovbWoPpL1CovzQr6OA). The scripts used to run the analyses and create the figures are available at https://github.com/rancilhac/Pogoniulus-phylogenomics-manuscript-data/tree/main.

