## Supplementary tables and figures for "Introgression across narrow contact zones shapes the genomic landscape of phylogenetic variation in an African bird clade"

**Table S1.** Samples included in the WGS dataset. Source of sample given if not from fieldwork performed by UCY. Source abbreviations: ZMUC=Zoological Museum of University of Copenhagen, LSUMNS - Louisiana State University Museum of Natural Science, CTR - UCLA Center for Tropical Research. The last column indicates the composition of three sample subsets used for phylogenetic inferences.

| **Species** | **Sample / Source** | **Locality** | **Sex** | **Subsets** |
| --- | --- | --- | --- | --- |
| *P. p. affinis* | AS32044 | Dolo Mena, Ethiopia | Female | Contact zones, Control |
| *P. p. affinis* | AS63208 | Greek River, Uganda | Male | Contact zones, Control |
| *P. p. affinis* | AS63245 | Grumeti, Tanzania | Female | Contact zones |
| *P. p. affinis* | AS63271 | Ruaha, Tanzania | Male | Contact zones, Control |
| *P. p. affinis* | J31315 | Diani, Kenya | Male | Core range |
| *P. p. affinis* | J31326 | Ruma, Kenya | Male | Contact zones, Control |
| *P. p. affinis* | J31330 | Sabaki, Kenya | Male | Core range |
| *P. p. affinis* | J31340 | Kisamisi, Kenya | Male | Contact zones |
| *P. p. affinis* | K38705 | Watamu, Kenya | Male | Core range |
| *P. p. affinis* | t132482,  ZMUC | Marsabit, Kenya | Male | Contact zones, Control |
| *P. c. chrysoconus* | AS39373 | Pian Upe, Uganda | Male | Contact zones, Control |
| *P. c. chrysoconus* | AS53106 | Acha, Uganda | Female | Contact zones, Control |
| *P. c. chrysoconus* | B39269, LSUMNS | Ghana | Male | Core range |
| *P. c. chrysoconus* | B39310, LSUMNS | Ghana | Female | Core range |
| *P. c. chrysoconus* | B39538, LSUMNS | Ghana | Male | Core range |
| *P. c. chrysoconus* | J31320 | Songhor, Kenya | Male | Contact zones, Control |
| *P. c. chrysoconus* | t1010216,  CTR | Malape, Cameroon | Male | Core range |
| *P. c. extoni* | AS32032 | Buffelskloof, South Africa | Male | Contact zones, Control |
| *P. c. extoni* | AS39210 | Chobe enclave, Botswana | Female | Core range |
| *P. c. extoni* | AS39211 | Chobe enclave, Botswana | Male | Core range |
| *P. c. extoni* | AS39277 | Seringveld, South Africa | Male | Contact zones, Control |
| *P. c. extoni* | AS39284 | Rayton, South Africa | Male | Contact zones |
| *P. c. extoni* | AS63269 | Rungwa, Tanzania | Female | Contact zones |
| *P. c. extoni* | AS63282 | Kitungole, Tanzania | Female | Contact zones, Control |
| *P. c. extoni* | K69305 | Uvinza, Tanzania | Male | Contact zones |
| *P. c. extoni* | R2 | Seringveld, South Africa | Male | Contact zones, Control |
| *P. c. extoni* | R4 | Seringveld, South Africa | Male | Contact zones |
| *P. p. pusillus* | AR93139 | Lake Eland, South Africa | Male | Core range |
| *P. p. pusillus* | AS53181 | Shongweni, South Africa | Male | Core range |
| *P. p. pusillus* | AS53184 | Shongweni, South Africa | Male | Core range |
| *P. p. pusillus* | DCK1 | Doreen Clark, South Africa | Male | Contact zones, Control |
| *P. p. pusillus* | Isoseq | Shongweni, South Africa | Male | Core range |
| *P. p. pusillus* | R14 | Harold Johnson  South Africa | Male | Contact zones, Control |
| *P. p. pusillus* | R19 | Shongweni, South Africa | Male | Core range |
| *P. p. uropygialis* | AS32046 | Wondo Genet, Ethiopia | Female | Contact zones |
| *P. p. uropygialis* | AS32086 | Checheho, Ethiopia | Female | Contact zones |
| *P. p. uropygialis* | AS32088 | Lalibela, Ethiopia | Female | Contact zones |
| *P. p. uropygialis* | AS32089 | Awash NP, Ethiopia | Male | Contact zones |
| *P. c. xanthostictus* | AS32040 | Dolo Mena, Ethiopia | Male | Contact zones |
| *P. c. xanthostictus* | AS32059 | Menagesha, Ethiopia | Female | Contact zones |
| *P. c. xanthostictus* | AS32064 | Bahir Dar, Ethiopia | Female | Contact zones |
| *P. c. xanthostictus* | AS32065 | Bahir Dar, Ethiopia | Male | Contact zones |
| *P. c. xanthostictus* | AS32069 | Bahir Dar, Ethiopia | Female | Contact zones |
| **Outgroups** | | | | |
| *P. atroflavus* | AS53113 | Semliki NP, Uganda | Male | – |
| *P. bilineatus* | AS53115 | Semliki NP, Uganda | Male | – |

**Table S2**. The 15 rooted 4-taxa topologies considered in the Topology Weighting analysis. aff-uro=*affinis* + *uropygialis*; chryso-xantho=*chrysoconus* + *xanthostictus*.

| **Name** | **Topology** | **Category** |
| --- | --- | --- |
| T1 | (((aff-uro,chryso-xantho),extoni),pusillus); | Geography |
| T2 | (((aff-uro,chryso-xantho),pusillus),extoni); | Geography |
| T3 | ((aff-uro,chryso-xantho),(extoni,pusillus)); | Geography |
| T4 | (((aff-uro,extoni),chryso-xantho),pusillus); | Geography |
| T5 | (((aff-uro,extoni),pusillus),chryso-xantho); | Geography |
| T6 | ((aff-uro,extoni),(chryso-xantho,pusillus)); | Geography |
| T7 | (((aff-uro,pusillus),chryso-xantho),extoni); | Color (red) |
| T8 | (((aff-uro,pusillus),extoni),chryso-xantho); | Color (red) |
| T9 | ((aff-uro,pusillus),(chryso-xantho,extoni)); | Color (both) |
| T10 | (aff-uro,(chryso-xantho,(extoni,pusillus))); | Geography |
| T11 | (aff-uro,(extoni,(chryso-xantho,pusillus))); | Others |
| T12 | (aff-uro,(pusillus,(chryso-xantho,extoni))); | Color (yellow) |
| T13 | ((aff-uro,(chryso-xantho,extoni)),pusillus); | Color (yellow) |
| T14 | ((aff-uro,(chryso-xantho,pusillus)),extoni); | Others |
| T15 | (chryso-xantho,(aff-uro,(extoni,pusillus))); | Geography |

**Table S3**. Parameters of the clines fitted to ancestry proxies (Admixture Q values) in the *chrysoconus* v. *affinis* and *extoni* v. *pusillus* contact zones. Values between brackets give 95% confidence intervals around parameter estimates. Note: by convention, the left side of the cline correspond to *affinis* or *pusillus*, and the right side to *chrysoconus* or *extoni.*

| **Cline** | **Width (km)** | **Center (km)** | **Left tail** | | **Right tail** | |
| --- | --- | --- | --- | --- | --- | --- |
|  |  |  | 𝛿 | 𝜏 | 𝛿 | 𝜏 |
| *chrysoconus* v. *affinis* | 10.0 [6.8, 15.6] | 0.4 [-0.6, 1.9] | – | – | – | – |
| *extoni* v. *pusillus* | 4.3 [1.2, 14.2] | 0.1 [-1.6, 2.3] | 1.1 [0.3, 3.8] | 0 [0, 0.3] | 1.8 [0.3, 6.1] | 0 [0, 0.3] |

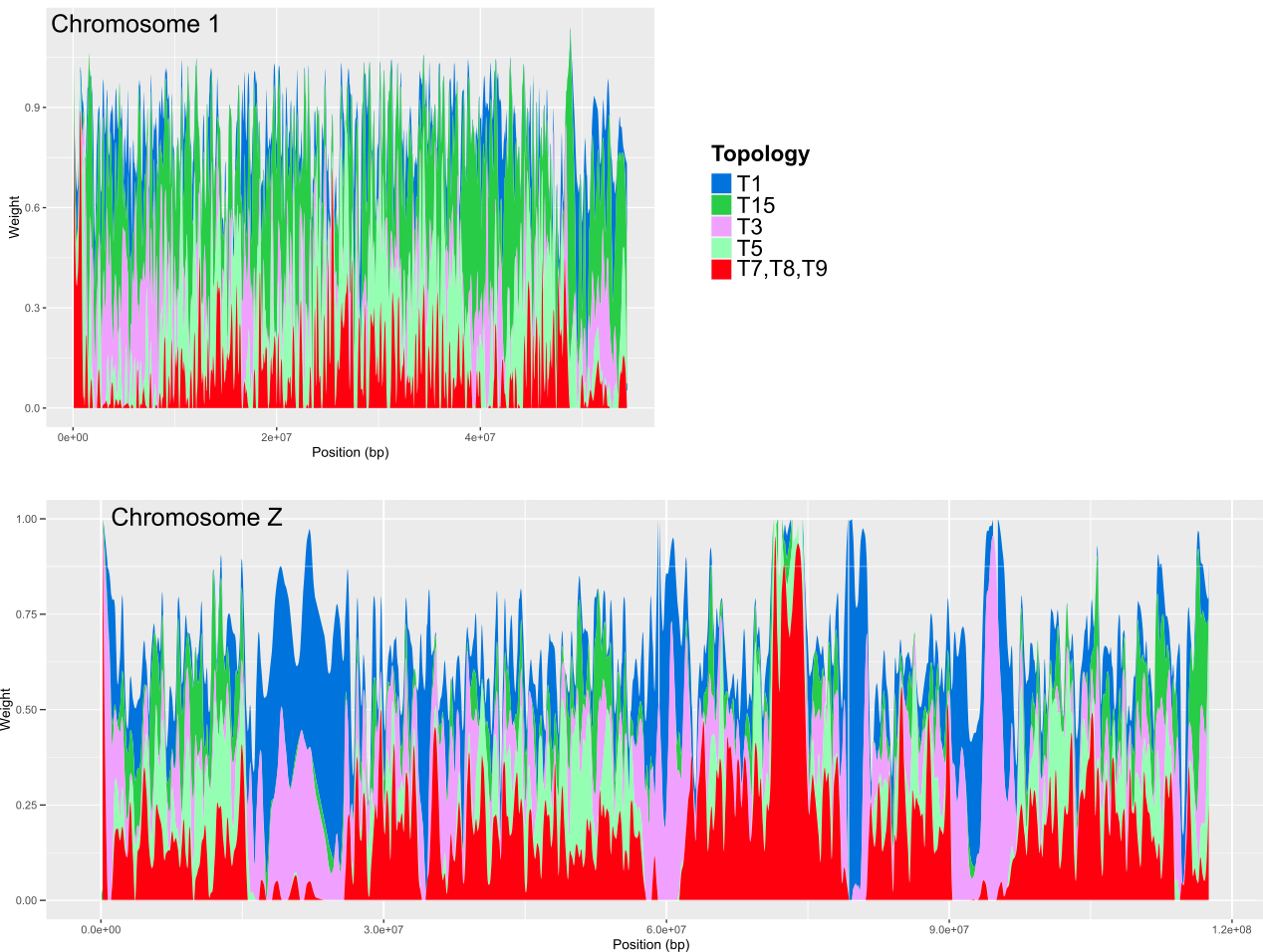

*.*

**Figure S1.** Distribution of the weights of the four commonest topologies (T1, T3, T5 and T15) as well as cumulative weights of the topologies where RFT are monophyletic (T7, T8 and T9) along chromosome 1 (top) and chromosome Z (bottom).

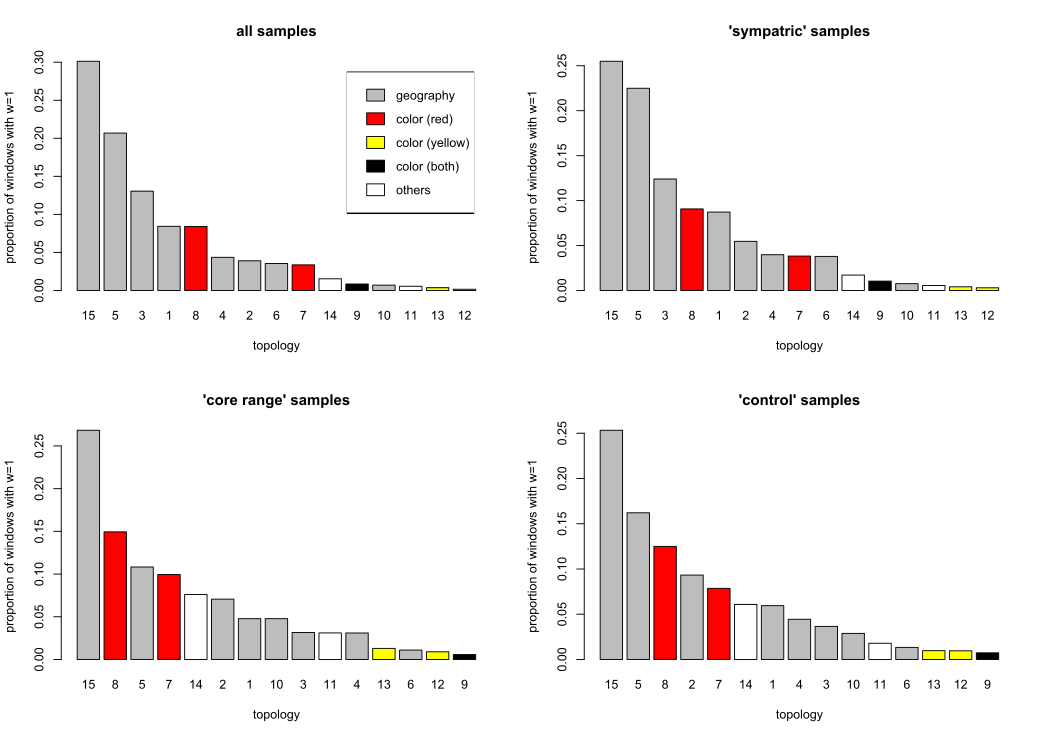

**Figure S2.** Proportion of autosomal 500 SNPs windows yielding full support (weight=1) to the 15 species tree topologies. The analysis was repeated on four sample sets: all samples; samples near contact zones; samples from core ranges; 14 random samples near the contact zones.The topologies are classified according to the species pairs supported as monophyletic (geography: species with a contact zone are monophyletic; color: either RFT, YFT or both are monophyletic; and others). The topologies are detailed in Table S1.

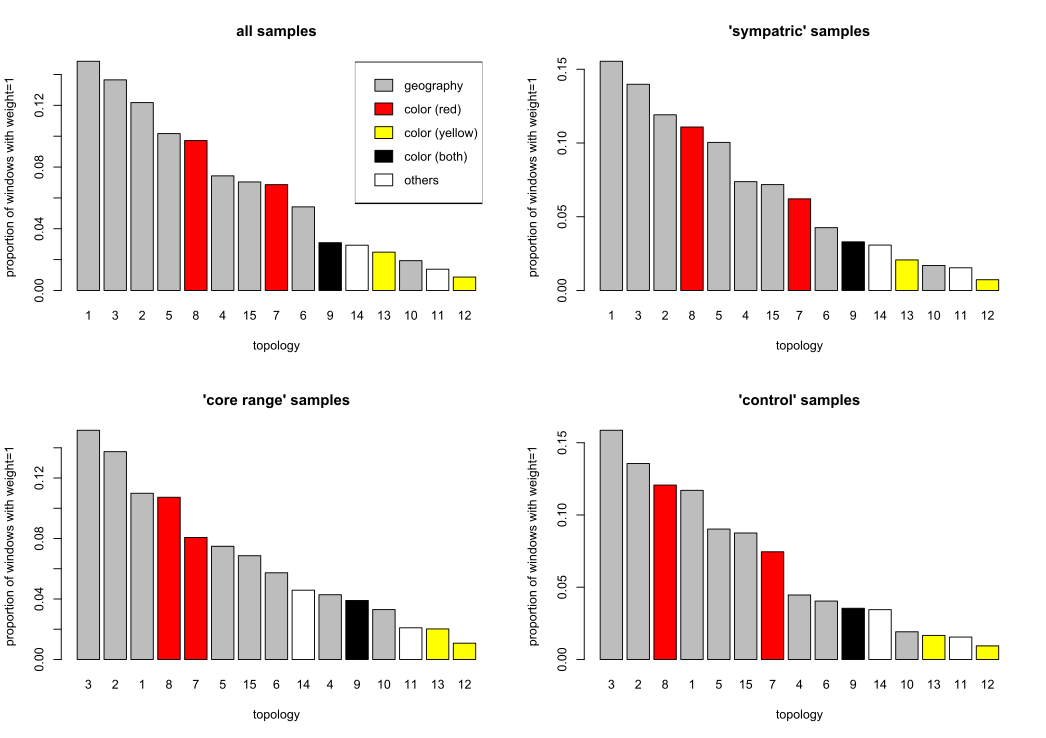

**Figure S3.** Proportion of 500 SNPs windows located on the Z chromosome yielding full support (weight=1) to the 15 species tree topologies. The analysis was repeated on four sample sets: all samples; near contact zones; samples from core ranges; 14 random samples near the contact zones.The topologies are classified according to the species pairs supported as monophyletic (geography: species with a contact zone are monophyletic; color: either RFT, YFT or both are monophyletic; and others). The topologies are detailed in Table S1.

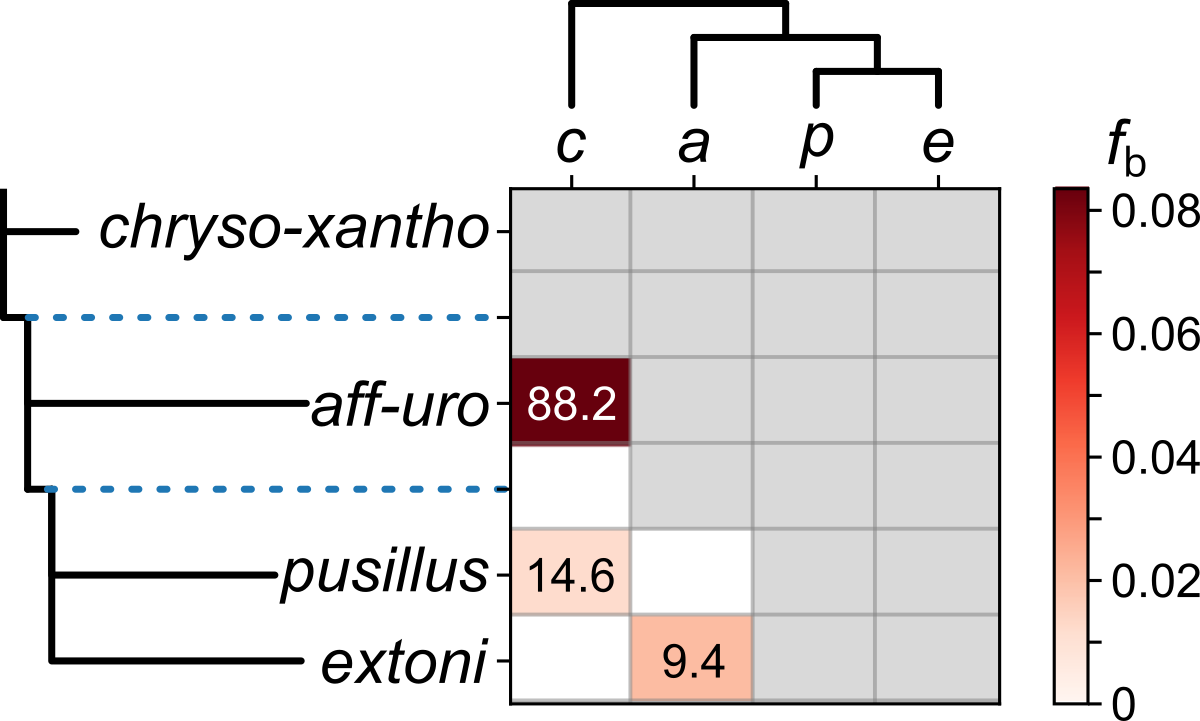

**Figure S4**. Fbranch matrix based on the dominant topology inferred with ASTRAL (T15). The numbers within the cells show the Z-score of the inferred introgression.

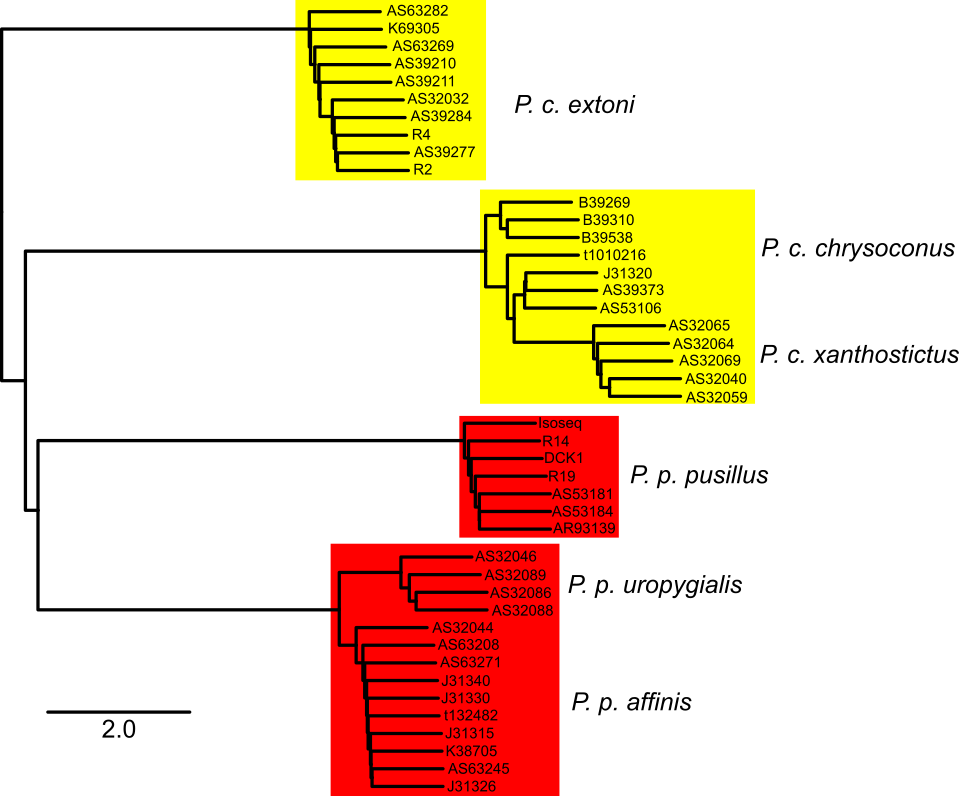

**Figure S5**. Phylogenetic tree inferred with ASTRAL based on 1,484 trees inferred from autosomal 500 SNPs sliding windows located in low recombination regions. All internal nodes received a posterior probability=1.0. The trees were rooted with sequences of *Pogoniulus atroflavus* and *P. bilineatus*, not represented for improved clarity. Branch lengths are expressed in coalescence units.

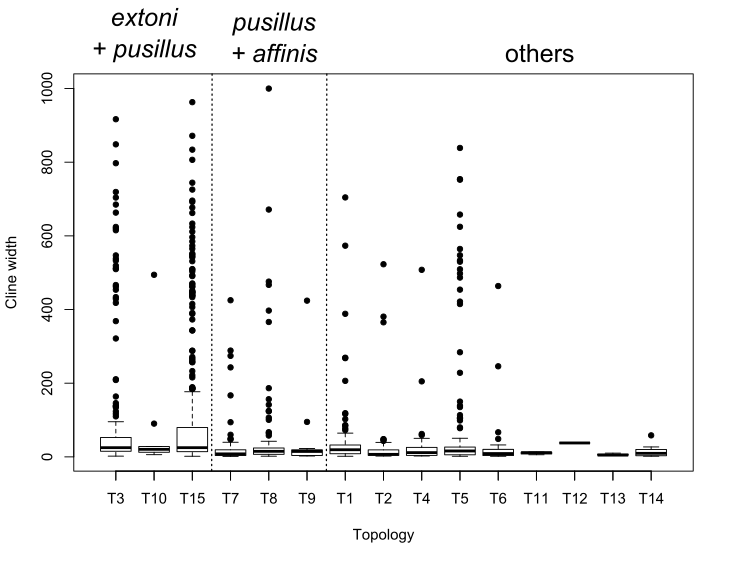

**Figure S6.** Distribution of cline width in the *extoni* v. *pusillus* contact zone for each of the 15 species tree topologies. Dots show outliers.

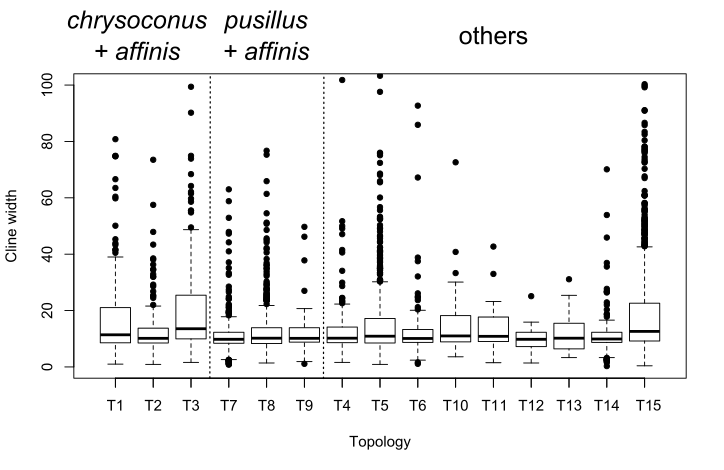

**Figure S7.** Distribution of cline width in the *chrysoconus* v. *affinis* contact zone for each of the 15 species tree topologies. Dots show outliers. The y-axis was shortened to 100 km for improved resolution.

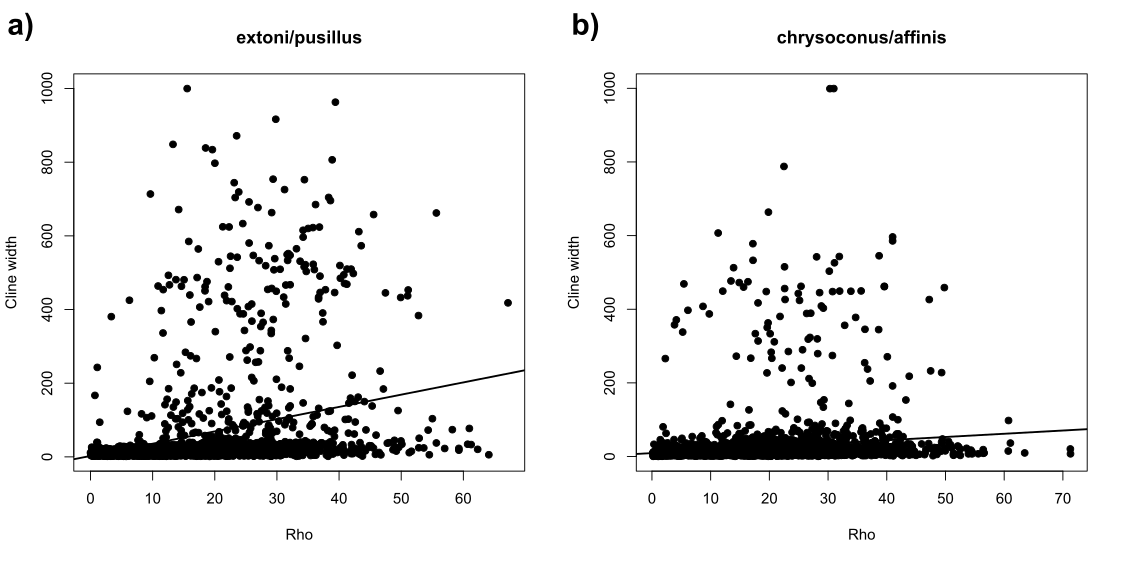

**Figure S8.** Correlation between cline width and population recombination rate (⍴) in a) the *extoni*/*pusillus* contact zone and b) the *chrysoconus*/*affinis* contact zone.

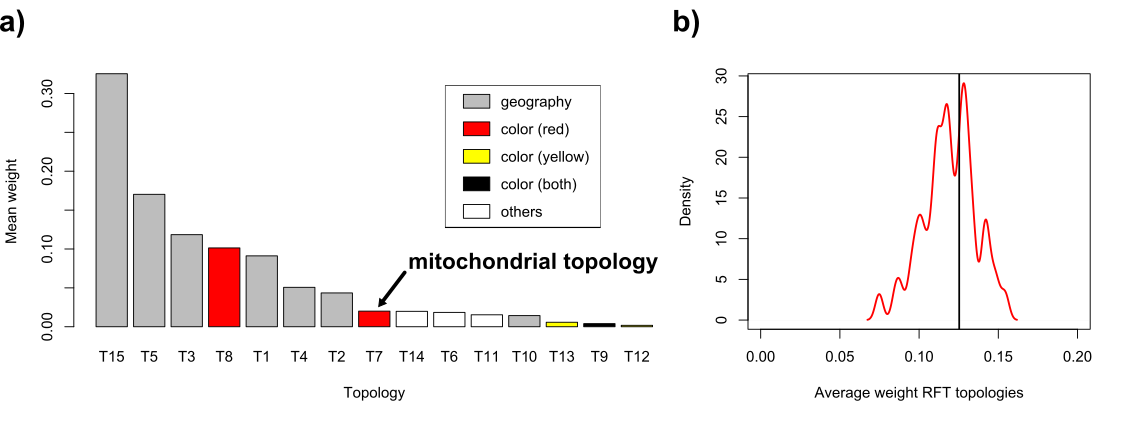

**Figure S9.** Phylogenetic signal around the coding sequences of mitochondria-interacting nuclear genes. a) Mean weights of the 15 topologies in 283 500-SNPs sliding windows overlapping these genes. The arrow indicates the topology supported by the mitochondrion; b) Density distribution of the combined average weights of T7,T8 and T9 (monophyletic RFT topologies) in 5,000 randomly drawn sets of 283 windows. The black vertical line shows the combined average weight in the windows overlapping the mitochondria-interacting genes.

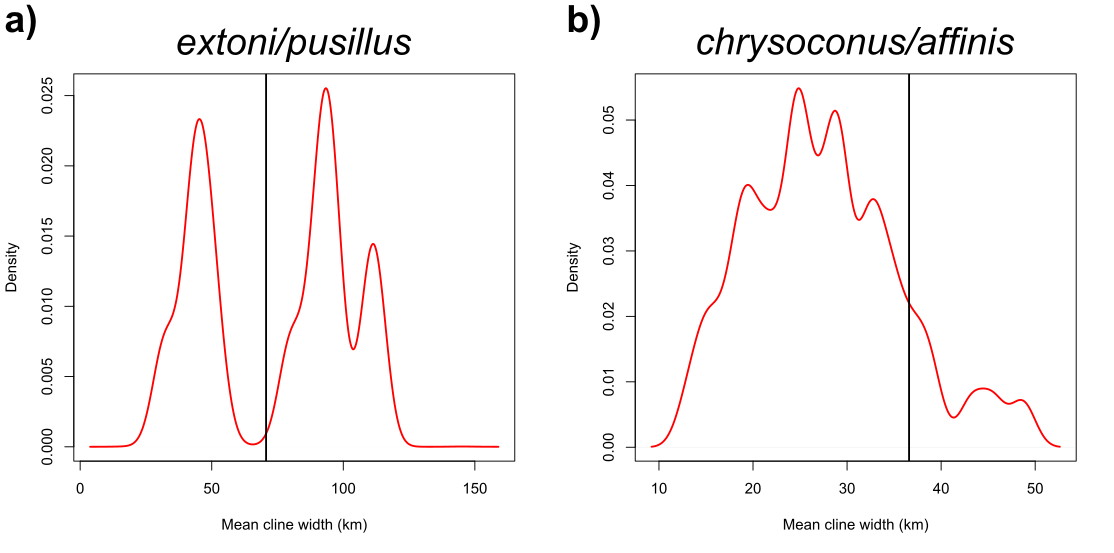

**Figure S10.** Cline width at SNPs near mitochondria-interacting nuclear genes. The plots show density distributions of mean cline width in 5,000 randomly drawn SNPs sets for a) the extoni/pusillus contact zone (sets of 38 SNPs) and b) the chrysoconus/affinis contact zone (sets of 80 SNPs). In both cases, the vertical black line shows the mean cline width for SNPs located near mitochondria-interacting nuclear genes.
